# An Autoimmunity-associated allele of *PTPN22* enhances innate antiviral immunity to protect against acute coronavirus infection

**DOI:** 10.1101/2025.07.15.664967

**Authors:** Alec M. Bevis, Kathryn J.L.H. Rosa, Nancy Schwarting, Tammy R. Cockerham, Catherine M. Kerr, Sunil More, Anthony R. Fehr, Robin C. Orozco

**Affiliations:** Department of Molecular Biosciences, University of Kansas; Lawrence, Kansas, 66045; Department of Veterinary Pathobiology, Oklahoma State University; Stillwater, Oklahoma, 74078

**Author notes:** Corresponding author. Address: 1200 Sunnyside Ave, Room 8035, Lawrence, KS 66045.

**Keywords:** PTPN22, coronavirus, Natural Killer (NK) cells

## Abstract

Allelic variation can impact viral clearance and disease severity. Still, our understanding of the effect of the autoimmunity-associated allelic variant of *Ptpn22* (PEP-R619W) on antiviral immunity remains incomplete, as previous reports have only focused on chronic Lymphocytic Choriomeningitis virus (LCMV) infection. This research defines how the loss of *Ptpn22* (PEP-null) and PEP-R619W changes antiviral immunity during an acute coronavirus infection. We address the hypothesis that CRISPR/Cas9-generated PEP-null and PEP-R619W mice have enhanced antiviral immunity over PEP-WT mice during coronavirus infection. Following Mouse Hepatitis Virus (MHV) A59 infection, we interrogated pathology, cytokine production, and cellular responses in the blood, spleen, and liver of PEP-WT, PEP-null, and PEP-R619W mice. Key findings show that PEP-R619W mice have reduced viral titer and weight loss, increased survival, and more mature natural killer (NK) cells in the liver and spleen compared to PEP-WT mice. Interestingly, protection against disease in PEP-null mice was inoculation-dose-dependent, whereas PEP-R619W conferred immunity regardless of infection dose. Further, Rag1-/- PEP-R619W mice had increased survival and reduced viral titer over Rag1-/- PEP-WT mice. PEP-R619W mice also had higher concentrations of IFNγ and enhanced IFNγ production by mature NK cells in the liver at 3 days post-infection. Finally, NK cell depletion elevated PEP-R619W viral titer to similar levels as PEP-WT mice. This is one of the first studies investigating the role of *Ptpn22* within NK cells and demonstrates that the *Ptpn22* allelic variant augments NK cell function and is beneficial during coronavirus infection.

**Significance Statement:** Approximately 5-15% of the North American population has the *PTPN22* 1858C>T allele, which has been linked with numerous autoimmune diseases and is considered the highest non-HLA risk allele for autoimmunity. Due to this link, the *PTPN22* 1858C>T allele is often considered pathologic or detrimental. However, recent studies have demonstrated the benefit of this allele in protection from chronic virus infection and some cancers. Yet, a significant research gap remains in understanding how the *PTPN22* 1858C>T allele impacts the immune response during acute, moribund infections. Using mice that are homozygous for the equivalent, autoimmunity-associated allele in *Ptpn22*, we demonstrate that this allele uniquely augments innate immunity and enhances Natural Killer cell function to protect against coronavirus infection.

## INTRODUCTION

Allelic variation in immune-regulatory genes is associated with protective or pathogenic immune responses during disease (1, 2). An example is *PTPN22* and its minor 1858C>T allele (rs2476601) (1–10). This *PTPN22* minor allele is strongly associated with multiple autoimmune diseases, has been cited as the highest non-HLA risk allele for autoimmunity, and is present in 5-15% of North Americans (1–10). The generational persistence of this *PTPN22* allele, despite its strong association with various severe autoimmune diseases, poses an evolutionary paradox. A theory is that the heightened immune responses it drives, though detrimental in autoimmunity, may offer protection against infections and cancer.

*PTPN22* encodes for the protein Lyp in humans and its ortholog PEP in mice. In humans and mice, *PTPN22* is expressed constitutively and exclusively within bone marrow-derived cells, which include all immune cells (3, 4). The *PTPN22* 1858C>T allele results in an amino acid substitution of an arginine (R) to tryptophan (W) that alters immune-regulatory function in both human (Lyp-R620W) and murine ortholog (PEP-R619W) models (3–6, 9, 11–21). To better understand the function of *Ptpn22*, prior studies have frequently employed *Ptpn22* knockout (PEP-null) mice. The molecular function of PEP has been best characterized as a negative regulator of T-cell receptor (TCR) and B-cell receptor (BCR) signaling, but its role in other immune cells is less well understood (3, 4, 11, 13, 16, 22–27). Further, the Lyp-R620W and PEP-R619W mutations do not directly impact phosphatase activity but lead to a loss of binding capacity with other proteins, such as Csk, TRAF3, Vav, Grb2, SKAP-HOM, and NLRP3, resulting in altered immune function (3, 4, 11–13, 17, 20, 21, 24, 28). Studies have defined several mechanisms in which the *PTPN22* 1858C>T allele potentiates heightened immune responses (3, 4, 11, 13, 16–21, 23, 24, 29). These same mechanisms that are detrimental during autoimmunity, such as enhanced T cell function, are protective when combating chronic viral infection and cancer (18, 19).

Previous studies show that both PEP-null and PEP-R619W mice, unlike wildtype (PEP-WT) mice, can clear chronic lymphocytic choriomeningitis virus clone 13 (LCMV-cl13) infection, and this phenotype is attributed to their enhanced antiviral T cell effector function (19, 30–32). However, the role of PEP and its PEP-R619W variant in virus infection has mostly been studied in the context of LCMV infection and T cells (19, 27, 30–33). Consequently, much remains unknown regarding the impact of PEP and its PEP-R619W variant on innate antiviral immunity, as well as whether PEP-null and PEP-R619W mice will exhibit similar immune responses during non-LCMV infection models. Despite the comparable immune phenotypes of PEP-null and PEP-R619W mice in LCMV-cl13 infection, other studies have highlighted several cellular and molecular differences between these two strains. For example, naïve PEP-null mice have an increased amount of Tregs, whereas PEP-R619W do not, compared to PEP-WT mice (26). Additionally, NLRP3-mediated IL-1β production is increased in PEP-R619W myeloid cells, but decreased in PEP-null myeloid cells, compared to PEP-WT cells (20, 21). These findings provide additional rationale to investigate differential responses between these strains, specifically within the context of viral infection. Therefore, we investigated whether PEP-null mice, PEP-R619W mice, or both can modulate innate antiviral immunity to confer protection during acute coronavirus infection.

In this study, we investigated the impact of *Ptpn22* (PEP) and its *Ptpn22* 1858C>T allele (PEP-R619W variant) on the innate immune response following acute coronavirus infection using the moribund, highly liver-tropic, mouse hepatitis virus (MHV) strain A59. Our findings show that PEP-R619W enhances natural killer (NK) cell maturation and strengthens NK effector functions, and that NK cells are necessary to restrict viral replication in the liver during innate time points in PEP-R619W mice. The results of this research define new mechanisms by which the autoimmunity-associated allele of *PTPN22* augments innate antiviral immunity and enhances protection against coronavirus infection.

## RESULTS

### PEP-R619W reduces disease and viral titer during MHV A59 infection

To determine the impact of *Ptpn22* and its autoimmunity-associated minor allele (PEP-R619W) during acute coronavirus infection, we infected wildtype (PEP-WT) and CRISPR-Cas9 generated *Ptpn22* knockout (PEP-null) and PEP-R619W mice with a common murine strain of coronavirus, mouse hepatitis virus (MHV) strain A59, which primarily infects the liver. In accordance with previous studies in the literature, MHV A59 infection did not cause severe disease following infection in 8-week-old mice, regardless of PEP-genotype (Fig. S1A and 1B) (34–40). Following 5e4 PFU MHV A59 infection, 5-week-old PEP-R619W and PEP-null mice had significantly reduced weight loss at day 5 post-infection (dpi) and 3-5 dpi, respectively, and this was paired with enhanced survival compared to PEP-WT mice (Fig. 1A and 1C). We next examined if the protection observed in PEP-null and PEP-R619W mice would persist at a higher inoculation dose, as early studies utilizing MHV A59 noted that higher viral input doses lead to more severe clinical symptoms (40). Following 1e5 PFU MHV A59 infection, PEP-R619W mice had significantly reduced weight loss at 3-6 dpi and enhanced survival compared to both PEP-WT and PEP-null mice (Fig. 1B and 1D). These data indicate that during 1e5 PFU infection, PEP-null mice are no longer protected from MHV A59 disease and instead phenocopy PEP-WT mice in weight loss and survival (Fig. 1A-D). To study the replication kinetics of MHV A59 *in vivo*, we quantified the viral load of the whole liver following 5e2 PFU infection, as it is the primary site of replication for MHV A59, at 1-, 3-, 5-, and 7-dpi via plaque assay. This dose was chosen as it has similar replication kinetics to higher inoculation doses, but does not induce severe clinical symptoms of weight loss and death, allowing for titer quantification that is not influenced by differences in survival at later timepoints (41, 42). PEP-R619W mice had significantly reduced viral titer at 1-, 3-, and 5-dpi compared to PEP-WT mice (Fig. 1E). However, PEP-null mice only demonstrated reduced titer at 5 dpi (Fig. 1F). Additionally, both PEP-R619W and PEP-null mice had accelerated clearance compared to PEP-WT mice, as 7/7 PEP-R619W mice and 7/8 PEP-null mice had cleared the infection by 7 dpi, whereas only 2/7 PEP-WT mice had cleared the infection at this time point (Fig. 1E and 1F). Splenic titers showed a low level of infection at 1-, 3-, and 5-dpi, regardless of genotype (Fig. S1C and S1D). All mice were able to clear the MHV A59 infection from the spleen by 7 dpi (Fig. S1C and S1D).

**Figure 1:**
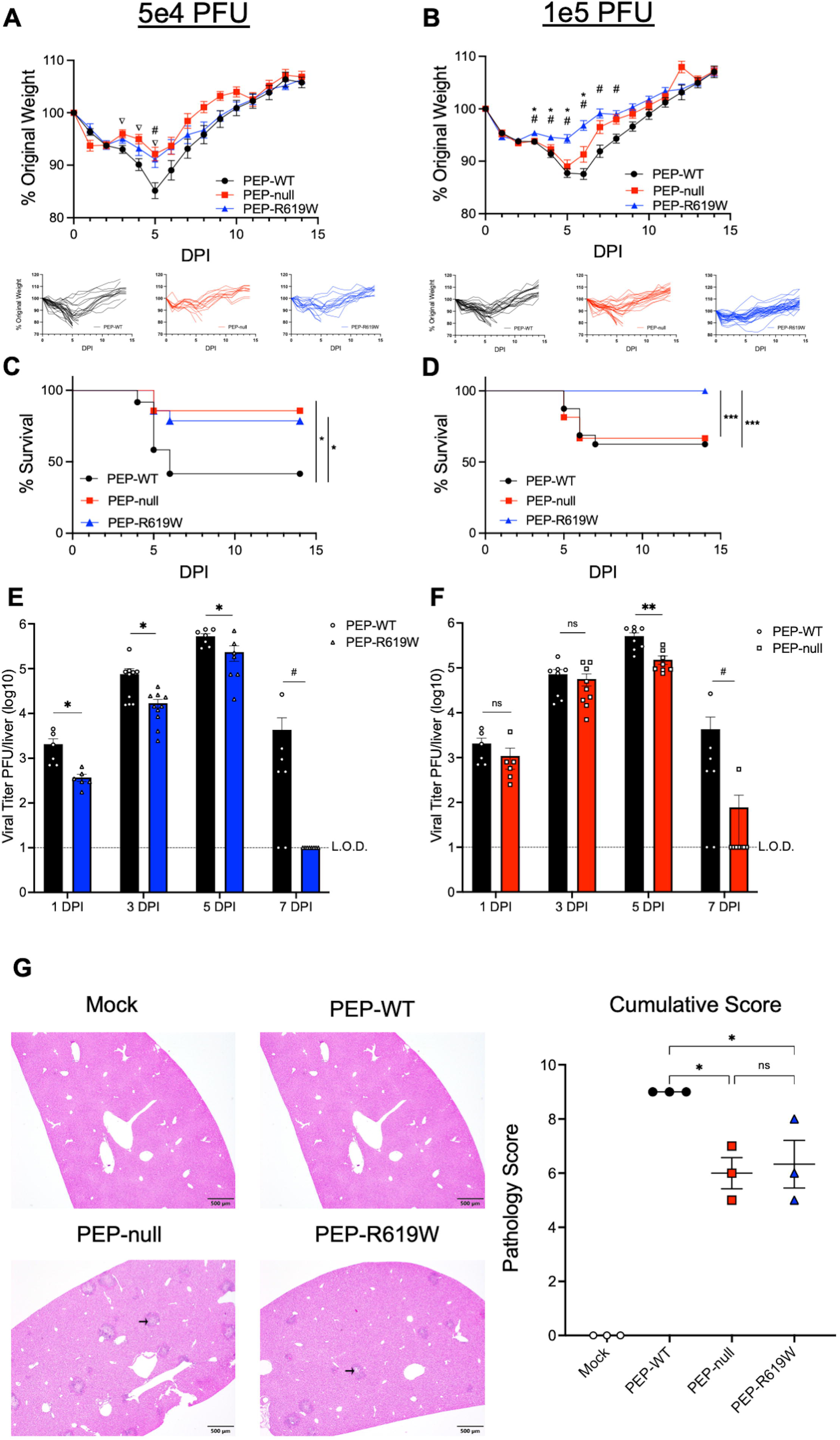
PEP-R619W mice have reduced disease during MHV A59 Infection. 5-week-old C57BL/6J mice with *Ptpn22* wild type (PEP-WT, black circle), *Ptpn22* knockout (PEP-null, red square), or the *Ptpn22* autoimmune-associated allele (PEP-R619W, blue triangle), were infected with 5e4 PFU (A, C) or 1e5 PFU (B, D) MHV A59 (i.p). Mice are tracked by weight (A, B) and survival (C, D). Respective spider plots for individual mouse weights are shown below (A, B). To measure viral burden, liver homogenates from infected (5e2 PFU) PEP-WT, PEP-null, and PEP-R619W mice were used for plaque assay (E, F). Histopathology of the livers of mice at 3 dpi following PBS injection (Mock) or 1e5 PFU MHV A59 infection, with representative photomicrographs of liver sections exhibiting areas of necrosis, inflammation, and edema/fibrin accumulation (black arrow) and cumulative pathology scores represented as mean + SEM. Scale bar = 500 μm (G). Weight loss and survival studies pooled from 3 individual experiments for each dose: Group sizes for 5e4 PFU infection (A, C) were PEP-WT n = 28, PEP-null n = 14, PEP-R619W n = 17. Group sizes for 1e5 PFU infection (B, D) were PEP-WT n = 31, PEP-null n = 27, PEP-R619W n = 31. For weight loss statistical analysis, significant p-values from One-way ANOVA with Tukey’s Multiple Comparisons at each dpi are represented by #, comparing PEP-WT vs. PEP-R619W, * comparing PEP-null vs. PEP-R619W, or ∇ comparing PEP-WT vs. PEP-null. Log-rank Mantel-Cox Test (survival), *p<0.05, ***p<0.001. For whole liver titer (E, F), each symbol represents an individual mouse. The dashed line indicates the limit of detection (L.O.D.). Certain PEP-WT mice are represented in both graphs, as some experiments had all three genotypes, whereas others contained only PEP-WT vs PEP-null or PEP-R619W. Combined data from 2 independent experiments, multiple unpaired T-Tests, PEP-WT vs PEP-R619W, PEP-WT vs PEP-null, at each time point, *p<0.01, **p<0.01. 7 dpi was analyzed using Fisher’s Exact Test, Cleared vs Not Cleared, #p<0.05. For histopathological analysis (G), mock mice received sterile PBS injection only and are a mixture of all three genotypes. Each dot represents an individual mouse, with p-values from One-way ANOVA with Tukey’s Multiple Comparisons shown, *p<0.05.

To determine if the reduced viral titer observed in the livers of PEP-R619W and PEP-null mice (Fig. 1E and 1F) was due to PEP-driven myeloid cell-intrinsic changes in these mice, resulting in decreased viral replication, we differentiated bone-marrow-derived macrophages (BMMs) and bone-marrow-derived dendritic cells (BMDCs). We infected them with MHV A59 *ex vivo*, as they are both readily infected *in vivo* by MHV (Fig. S1E) (43–45). We did not detect any significant differences in viral titer across the three genotypes in either BMMs (Fig. S1F) or BMDCs (Fig. S1G), indicating that PEP-R619W and PEP-null do not mediate protection by directly impacting viral replication through cell-intrinsic mechanisms. Histopathologic analysis of liver sections at 3 dpi following 1e5 PFU infection demonstrated multifocal regions of marked hepatocellular necrosis in PEP-WT mice, characterized by infiltration of inflammatory cells and deposition of edema and fibrin (black arrow) (Fig. 1G and Fig. S1H). In comparison, PEP-R619W and PEP-null mice exhibited markedly reduced hepatic injury, with mild necrosis and inflammation (Fig. 1G and Fig. S1H). These combined results show that PEP-R619W mice have improved disease outcomes and reduced viral burden, which includes accelerated clearance, during MHV A59 infection.

### PEP-R619W prolongs the survival of infected mice in the absence of T and B cells

Due to the enhanced protection against disease conferred by PEP-R619W at early days post-infection (Fig. 1), we wanted to determine if a strengthened innate immune response was sufficient to fully protect these mice, or if T and B cells were necessary for survival during MHV A59 infection. To do this, we infected Rag1-/- PEP-WT, Rag1-/- PEP-null, and Rag1-/- PEP-R619W mice with 5e4 PFU or 1e5 PFU MHV A59. Like the immune-competent mice, Rag1-/- PEP-R619W mice had noticeably less weight loss throughout most of the early stages of the infection (Fig. 2A and 2B) and mice survived longer (Fig. 2C and 2D) compared to Rag1-/- PEP-WT mice at both 5e4 and 1e5 PFU MHV A59 infection. Rag1-/- PEP-null mice also have prolonged survival over Rag1-/- PEP-WT at 5e4 PFU infection (Fig. 2C). However, at 1e5 PFU infection, PEP-null mice no longer had prolonged survival (Fig. 2D). These data also show that T and B cells are necessary for overall survival, regardless of PEP-genotype or MHV A59 dose, as all mice on a Rag1-/- background eventually succumb to disease (Fig. 2C and 2D). Next, we measured viral load in the liver at 3- and 5-dpi following 5e2 PFU MHV A59 infection. Rag1-/- PEP-R619W mice had reduced viral burden compared to Rag1-/- PEP-WT mice (Fig. 2E). No difference was detected between Rag1-/- PEP-WT and Rag1-/- PEP-null mice liver titers (Fig. 2F). There was no difference detected in splenic viral titers at 3 and 5 dpi between Rag1-/- PEP-WT and Rag1-/- PEP-R619W mice or Rag1-/- PEP-WT and Rag1-/- PEP-null mice (Fig. S2). Overall, these data suggest that PEP-null enhances the innate immune response during 5e4 PFU infection in both immune competent and Rag1-/- models to improve survival compared to PEP-WT mice, and these results are comparable to PEP-R619W mice (Figs. 1 and 2). However, only PEP-R619W mice showed enhanced protection following 1e5 PFU infection, suggesting that this autoimmune-associated variant modifies the innate response through additional mechanisms compared to PEP-null, and this confers increased resistance against MHV A59-associated disease (Figs. 1 and 2).

**Figure 2:**
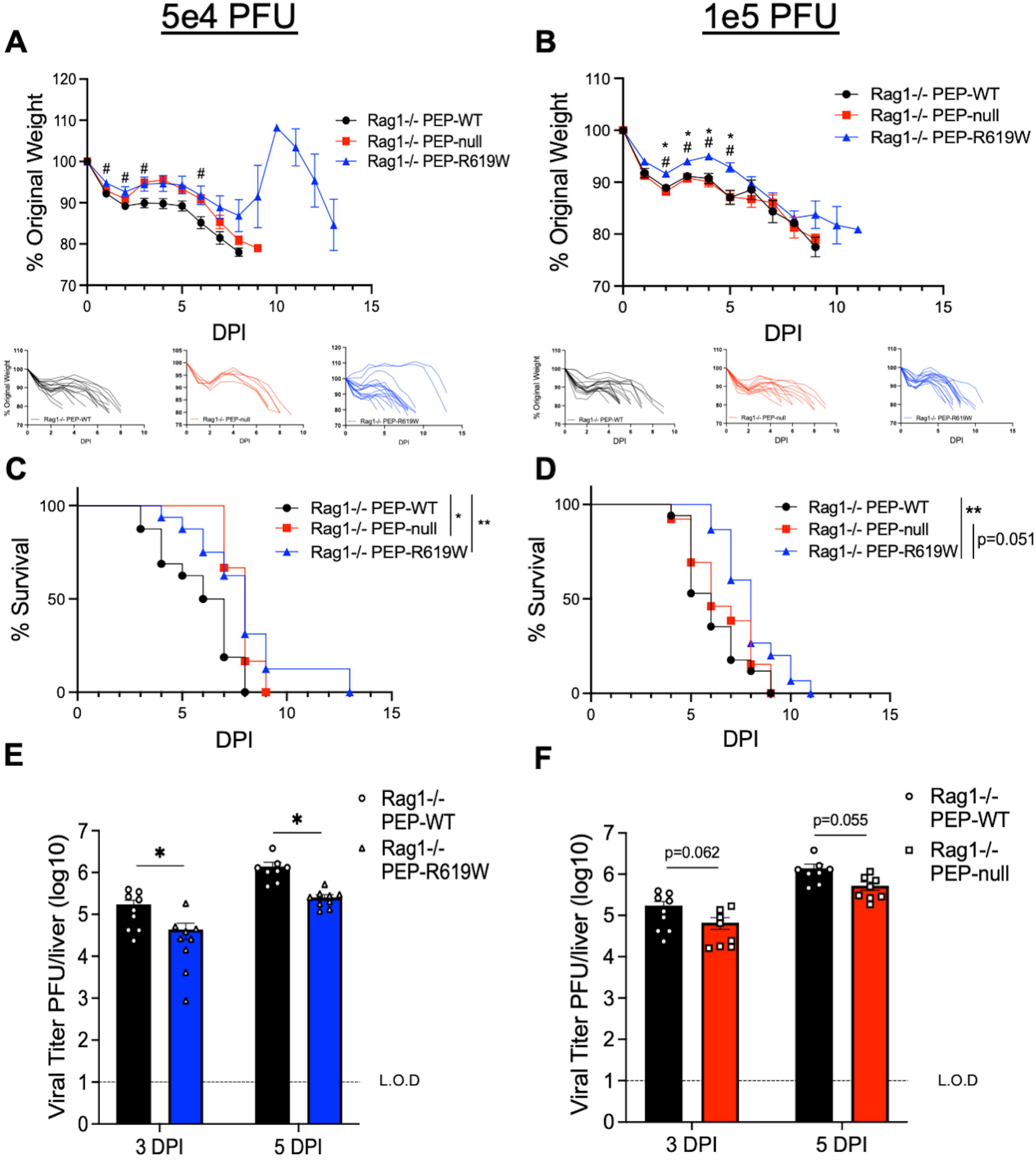
PEP-R619W innate cells provide some protection against disease, but T and B cells are necessary for survival. Rag1-/- PEP-WT (black circle), Rag1-/- PEP-null (red square), or Rag1-/- PEP-R619W (blue triangle), mice were infected with 5e4 PFU (A, C) or 1e5 PFU (B, D) MHV A59 (i.p), and were tracked by weight (A, B) and survival (C, D). Respective spider plots for individual mouse weights are shown below (A, B). To measure viral burden, liver homogenates from infected (5e2 PFU) Rag1-/- PEP-WT, Rag1-/- PEP-null, and Rag1-/- PEP-R619W mice were used for plaque assay (E, F). Weight loss and survival studies pooled from 2 individual experiments for each dose: Group sizes for 5e4 PFU (A, C) were Rag1-/- PEP-WT n = 16, Rag1-/- PEP-null n = 6, Rag1-/- PEP-R619W n = 16. Group sizes for 1e5 PFU (B, D) were Rag1-/- PEP-WT n = 17, Rag1-/- PEP-null n = 13, Rag1-/- PEP-R619W n = 15. For weight loss statistical analysis, significant p-values from One-way ANOVA with Tukey’s Multiple Comparisons at each dpi are represented by # comparing Rag1-/- PEP-WT vs. Rag1-/- PEP-R619W and * comparing Rag1-/- PEP-null vs. Rag1-/- PEP-R619W. Log-rank Mantel-Cox Test (survival), *p<0.05, **p<0.01. For whole liver titers (E, F), each symbol represents an individual mouse. The dashed line indicates the limit of detection (L.O.D.). Certain Rag1-/- PEP-WT mice are represented in both graphs, as some experiments had all three genotypes, whereas others contained only Rag1-/- PEP-WT vs Rag1-/- PEP-null or Rag1-/- PEP-R619W. Combined data from 2 independent experiments, multiple unpaired T-Tests, Rag1-/- PEP-WT vs Rag1-/- PEP-R619W, Rag1-/- PEP-WT vs Rag1-/- PEP-null, at each time point. *p<0.01, **p<0.01.

### PEP-R619W mice have increased innate immune presence in the liver and spleen during MHV A59 infection

We next aimed to determine how the PEP-R619W variant augmented the innate immune response. We focused on 3 dpi because 1) this is when we can consistently observe a divergence in weight loss between genotypes, 2) PEP-R619W mice have reduced liver titer and pathology at this time point, and 3) this is a time point at which the innate response dominates, whereas at 5 dpi, there could be some early T cell activity. We first investigated the innate cellular response in the spleen after 5e4 PFU MHV A59 infection, as we wanted to determine if the immune response in PEP-null and PEP-R619W mice would correlate with the improved weight loss and survival phenotypes observed at this dose. Overall, PEP-R619W and PEP-null mice had comparable immune phenotypes at 3 dpi in the spleen post-infection (Fig. S3). There was no difference detected in the bulk number of splenocytes (Fig. S3B) or frequency of CD3e- CD19-(innate cells) of all live cells (Fig. S3C), regardless of infection or PEP genotype. However, PEP-R619W and PEP-null mice had significantly increased frequencies of NK cells and DCs compared to infected PEP-WT mice (Fig. S3D).

Interestingly, PEP-R619W mice had more splenic CD11b+ NK cells than PEP-WT, indicating PEP-R619W NK cells are significantly more mature than PEP-WT NK cells following infection (Fig. S3E) (46, 47). Further, there were no differences detected in the frequency of CD8α+ conventional dendritic cells (CD8α+ cDCs), which are likely type 1 conventional dendritic cells (cDC1s) (Fig. S3F) (48). While the expression of activation markers CD80 and CD86 was significantly upregulated post-infection in splenic CD8α+ cDCs, no significant difference was detected between infected genotypes (Fig. S3G and S3H, respectively). Similar results were seen in splenic CD8α-cDCs, which are likely type 2 conventional dendritic cells (cDC2s) (Fig. S3I-K), though there was a decrease in CD86 expression in PEP-null and PEP-R619W splenic CD8α-cDCs compared to PEP-WT post-infection (Fig. S3K) (48).

Next, we wanted to know if PEP-R619W mice had an altered immune response in the liver, the main site of viral replication, during the 1e5 PFU MHV A59 infection, as they were the only genotype protected at the inoculation dose (Figs. 1 and 2). At 3 dpi, we did not detect significant differences in the absolute number of liver cells (Fig. 3A) or in the frequency (%) of CD45+ cells (Fig. 3B) between our mock mice (uninfected controls) and our infected PEP-WT, PEP-null, and PEP-R619W mice. However, PEP-R619W mice have a higher proportion of CD3e- CD19- of CD45+ immune cells, which we define as an immune “innate subset,” compared to PEP-WT mice post-infection (Fig. 3C). Delving further into this innate cell population, PEP-R619W mice had increased proportions of natural killer (NK) cells, dendritic cells (DCs), and macrophages of CD45+ cells over both PEP-WT and PEP-null mice post-infection (Fig. 3D). We did not detect any differences between infected genotypes in plasmacytoid DCs (pDCs), monocytes, or neutrophils (Fig. 3D). We also did not observe any differences in the proportion of or CD86 expression on Tim4+ macrophages, which are likely Kupffer cells amongst infected PEP-WT, PEP-null, and PEP-R619W mice (Fig. S4A-C) (49–51). Post-infection, all genotypes increased CD86 expression on Tim4+ macrophages (Fig. S4C). Within the DC population, there was a decrease in both the frequency and expression of MHC-II (Fig. S4D and S4E, respectively) in infected genotypes compared to uninfected controls (mock). Further, we did not identify any differences in the frequency and expression of CD86 on DCs between infected genotypes (Fig. S4F and S4G).

**Figure 3:**
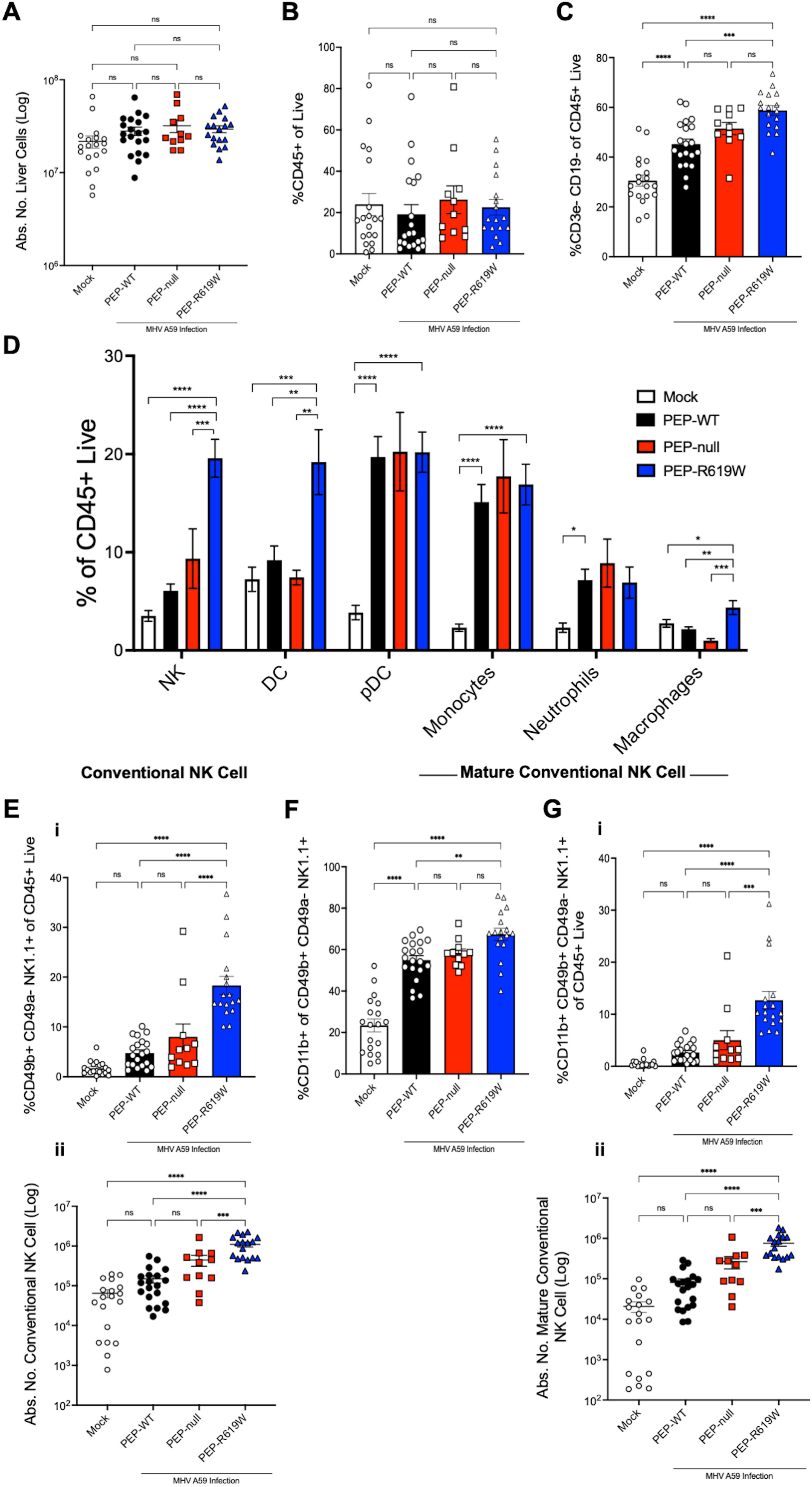
PEP-R619W mice have more innate immune and mature NK cells in the Liver. Following PBS injection (Mock, white bar, circle) or 1e5 PFU MHV A59 infection (i.p) in PEP-WT (black bar, circle), PEP-null (red bar, square), and PEP-R619W (blue bar, triangle) mice, at 3 dpi the liver was perfused, harvested, and antibody stained for flow cytometric analysis. The absolute count of liver cells for all genotypes (A). Frequencies (%) of CD45+ cells of live cells (B) and of CD3e- CD19- of CD45+ Live (C). Frequency (%) of several innate cells of CD45+ cells is shown (D). Gating strategy for all innate subsets is: Lymphocytes> Single cell x2> autofluorescent-> Live> CD45+> CD3e- CD19-> Ly6G- Ly6C+/- (non-neutrophils). NK Cells: innate subsets> NK1.1+. DCs: innate subsets> NK1.1-> CD11c+ PDCA-1-. pDCs: innate subsets> NK1.1-> PDCA-1+ CD11c+/-. Monocytes: innate subsets> NK1.1-> Ly6C^hi^ CD11b+. Macrophage: innate subsets> NK1.1-> Ly6C-CD11b+/-> F4/80+ CD11b+. Neutrophils: Lymphocytes> Single cell x2> autofluorescent-> Live> CD3e-CD19-> Ly6G+ Ly6C+. Frequency (%, i) of CD45+ and absolute count (#, ii) of conventional NK cells (E). Frequency (%) mature conventional NK cells (F). Frequency (%, i) of CD45+ and absolute count (#, ii) mature conventional NK cells (G). Gating strategy for NK subsets: Conventional NK Cells: innate subsets> NK1.1+> CD49b+ CD49a-. Mature Conventional NK Cells: Conventional NK Cells> CD11b+. Quantification of flow cytometric data was pooled from 6 independent experiments. Each symbol represents an individual mouse. Mock mice are a mixture of all 3 genotypes. p-values from One-way ANOVA with Tukey’s Multiple Comparisons are shown for each panel. *p<0.05, **p<0.01, ***p<0.001, ****p<0.0001.

MHV A59-infected PEP-R619W mice had more conventional NK cells compared to infected PEP-WT and PEP-null mice, characterized as CD49b+ CD49a- of NK1.1+ cells, whereas the inverse is considered tissue-resident type 1 innate lymphoid cells (ILC-1) (Fig. 3E) (52, 53). Additionally, PEP-R619W conventional NK cells were more mature (CD11b+) compared to PEP-WT (Fig. 3F), and overall, PEP-R619W mice had more mature conventional NK cells (CD11b+ CD49b+ CD49a-NK1.1+) compared to both PEP-WT and PEP-null mice during coronavirus infection (Fig. 3G). These mature NK cells are of particular interest because, as NK cells mature, they enhance their effector function potential, including increased interferon-gamma (IFNγ) production and elevated expression of cytotoxic molecules, such as perforin and granzyme B (46, 47). Collectively, these data demonstrate that the PEP-R619W variant enhances both splenic and liver innate antiviral immunity, specifically increasing the overall amount and maturity of NK cells.

### PEP-null splenic mature NK cell effector functions are changed depending on the MHV A59 inoculation dose

Since our data showed that PEP-R619W mice have significantly more mature NK cells in both the liver and spleen at 3 dpi, we next wanted to determine if the NK cells amongst our mice had changed functional properties, and if the inoculation dose of MHV A59 would impact these phenotypes. Again, we observed inoculation dose-dependent differences, where at 5e4 PFU, PEP-null and PEP-R619W mice had similar NK cell immune phenotypes in the spleen (Fig. S5A-F). Specifically, following 5e4 PFU infection, PEP-null and PEP-R619W mature splenic NK cells had less CD107a, a marker of degranulation (Fig. S5A and S5B), increased perforin (Fig. S5C and S5D), and no detectable difference in IFNγ (Fig. S5E and S5F) compared to PEP-WT mature splenic NK cells. Yet at 1e5 PFU, PEP-null and PEP-WT phenotypes were more similar in the spleen (Fig. S5G-L). With 1e5 PFU infection, PEP-R619W mature splenic NK cells had reduced degranulation marker CD107a (Fig. S5G and S5H), increased perforin (Fig. S5I and S5J), and an increase in frequency, but not expression, of IFNγ (Fig. S5K and S5L) compared to PEP-WT and PEP-null mature splenic NK cells. These data provide some insight into why PEP-null mice have differing phenotypes (survival) during different doses of MHV A59 infection, as differential immune activation at 1e5 PFU infection may be responsible for the previously described increase in morbidity (Figs. 1 and 2).

### PEP-R619W enhances liver mature NK cell effector function and the overall production of IFN**γ** during virus infection

As only PEP-R619W mice, but not PEP-null, had more mature conventional NK (M-cNK) cells compared to PEP-WT mice at 3 dpi in the liver, we next investigated if PEP-R619W M-cNK cells have altered effector functions compared to PEP-WT M-cNK cells. Following 1e5 PFU infection, at 3 dpi in the liver, PEP-R619W M-cNK cells (CD11b+ CD49b+ CD49a-NK1.1+) have higher expression of (i) IFNγ, (ii) T-bet, (iii) granzyme B, and (iv) perforin, compared to PEP-WT M-cNK cells (Fig. 4A, i-iv). There was no difference detected in the expression of (v) TNFα or (vi) CD107a between PEP-WT and PEP-R619W M-cNK cells (Fig. 4A, v-vi). Further, PEP-R619W M-cNK cells were more poly-functional compared to PEP-WT, as there was an increased proportion of IFNγ+ Perforin+ and IFNγ+ Perforin+ TNFα+ M-cNK cells in PEP-R619W mice.

**Figure 4:**
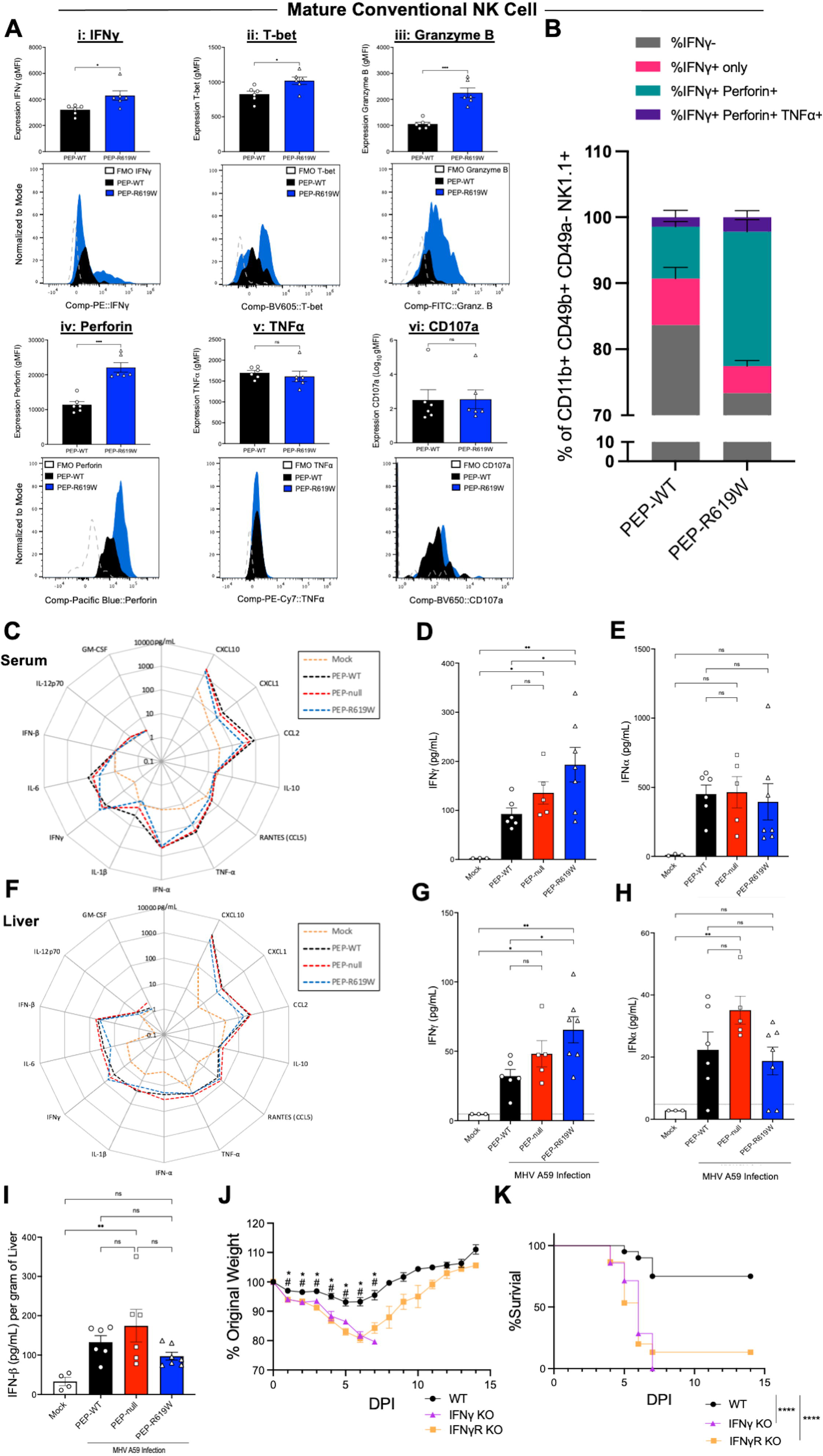
PEP-R619W augments mature conventional NK antiviral functions during coronavirus infection. 3 dpi liver M-cNK analysis following 1e5 PFU MHV A59 infection (i.p) in PEP-WT (black bar, circle) and PEP-R619W (blue bar, triangle) mice. Expression (gMFI) of IFNγ (i), T-bet (ii), Granzyme B (iii), Perforin (iv), TNFα (v), and CD107a (vi) in Mature Conventional NK cells, with representative histograms, with fluorescence minus one (FMO) controls included for each marker, below (A). Proportion of IFNγ-(grey), IFNγ+ only (pink), IFNγ+ Perforin+ (teal), and IFNγ+ Perforin+ TNFα+ (purple) of Mature Conventional NK Cells (B). Expression of 13 different cytokines and chemokines were quantified using LEGENDplex. Radar plots showing the 13 different cytokines from the serum (C) and liver (F), with select cytokines, IFNγ (D, G) and IFNα (E, H), quantified to the right. IFNβ concentration in the liver at 3 dpi [ELISA] (I). C57BL/6 mice with wild type (PEP-WT, black circle), IFNγ knockout (IFNγ KO, purple triangle), or IFNγ Receptor knockout (IFNγR KO, orange square) were infected with 5e4 PFU MHV A59 (i.p). Mice are tracked by weight (J) and survival (K). Quantification of flow cytometric data is representative of 3 independent experiments. p-values from Welch’s unpaired t test are shown, *p<0.05, **p<0.01, ***p<0.001. LEGENDplex assays are from 1 experiment. Each symbol represents an individual mouse. Mock mice received PBS injections only and are a mixture of all 3 genotypes. p-values from One-way ANOVA with Tukey’s Multiple Comparisons are shown for each panel. *p<0.05, **p<0.01, ***p<0.001. Weight loss and survival studies pooled from 2 individual experiments. Group sizes were PEP-WT n = 19, IFNγ KO n = 4, and IFNγR KO n = 15. For weight loss statistical analysis, significant p-values from One-way ANOVA with Tukey’s Multiple Comparisons at each dpi are represented by # comparing PEP-WT vs. IFNγ KO, or * PEP-WT vs. IFNγR KO. Log-rank Mantel-Cox Test (survival), ****p<0.0001.

Next, we examined the magnitude of the cytokine/chemokine response both systemically and in the whole liver at 3 dpi. Strikingly, out of the thirteen different cytokines and chemokines paneled in the serum (Fig. 4C), only the concentration of IFNγ was significantly increased in PEP-R619W mice (Fig. 4D), yet all other molecules, including interferon-alpha (IFNα) (Fig. 4E), which is necessary for protection against MHV A59 infection, were unchanged between infected genotypes (54). Similar results were detected in the whole liver homogenate (Fig. 4F-H). To corroborate these findings, we measured the levels of IFNβ in the liver at 3 dpi in our PEP-WT, PEP-null, and PEP-R619W mice. We detected no difference between infected genotypes (Fig. 4I).

Given the enhanced M-cNK IFNγ expression and the overall increased IFNγ concentration in our PEP-R619W mice, we next wanted to determine if IFNγ and/or IFNγ-signaling was necessary for survival and weight retention during MHV A59 infection. To test this, we infected WT, IFNγ knockout (IFNγ KO), and IFNγ receptor knockout (IFNγR KO) mice with 5e4 PFU of MHV A59. Both IFNγ KO and IFNγR KO mice had more severe weight loss (Fig. 4J) and nearly 100% mortality (Fig. 4K) compared to WT mice, demonstrating the importance of IFNγ for protection during MHV A59 infection.

Taken together, these data show that PEP-R619W liver M-cNK cells produce more IFNγ and are more poly-functional than PEP-WT M-cNK cells, contributing to the increased levels of IFNγ in both the serum and liver at 3 dpi. Further, IFNγ and IFNγ signaling are critical for survival. This suggests that the protection conferred by PEP-R619W during MHV A59 infection may be driven, in part, by enhanced NK cell IFNγ production.

### NK Cell maturation is accelerated in PEP-null and PEP-R619W mice

Due to the increased presence of mature NK cells in the spleens (CD11b+ of NK1.1+) and liver (CD11b+ of CD49b+ CD49a-NK1.1+) of PEP-R619W mice post-infection (Fig. S3E, Fig. 3E-G), we examined if PEP-R619W NK cells had differential maturation states compared to PEP-WT at homeostasis. Previous studies have shown that murine NK cells follow a four-stage developmental program, defined by the subsequent acquisition and loss of surface receptors CD27 and CD11b, and these maturation states can be seen in the spleen (46, 47). The four stages, from least to most mature, are defined as; (i) CD27- CD11b- (Double Negative, DN), (ii) CD27+ CD11b-, (iii) CD27+ CD11b+ (Double Positive, DP), and (iv) CD27-CD11b+ (46, 47). This developmental program is associated with the stepwise acquisition of NK cell effector function, and the accumulation of terminally mature NK cells (CD27-CD11b+) plateaus at 8 weeks of age (46, 47). At 5 weeks of age, PEP-null and PEP-R619W splenic NK cells had differential maturation states compared to PEP-WT (Fig. 5). Specifically, of the NK cell repertoire, PEP-WT mice had more DP NK cells, but PEP-null and PEP-R619W had increased proportions of terminally mature (CD27-CD11b+) NK cells at 5 weeks of age (Fig. 5A, 5D, 5E). However, at 8 weeks of age, there were no differences detected in the proportion of DP or terminally mature NK cells between the three genotypes of mice. Collectively, these data indicate that PEP may negatively impact NK cell maturation at homeostasis, which could play a role in the observed differences in mature NK cell presence and function in PEP-R619W mice following MHV A59 infection.

**Figure 5:**
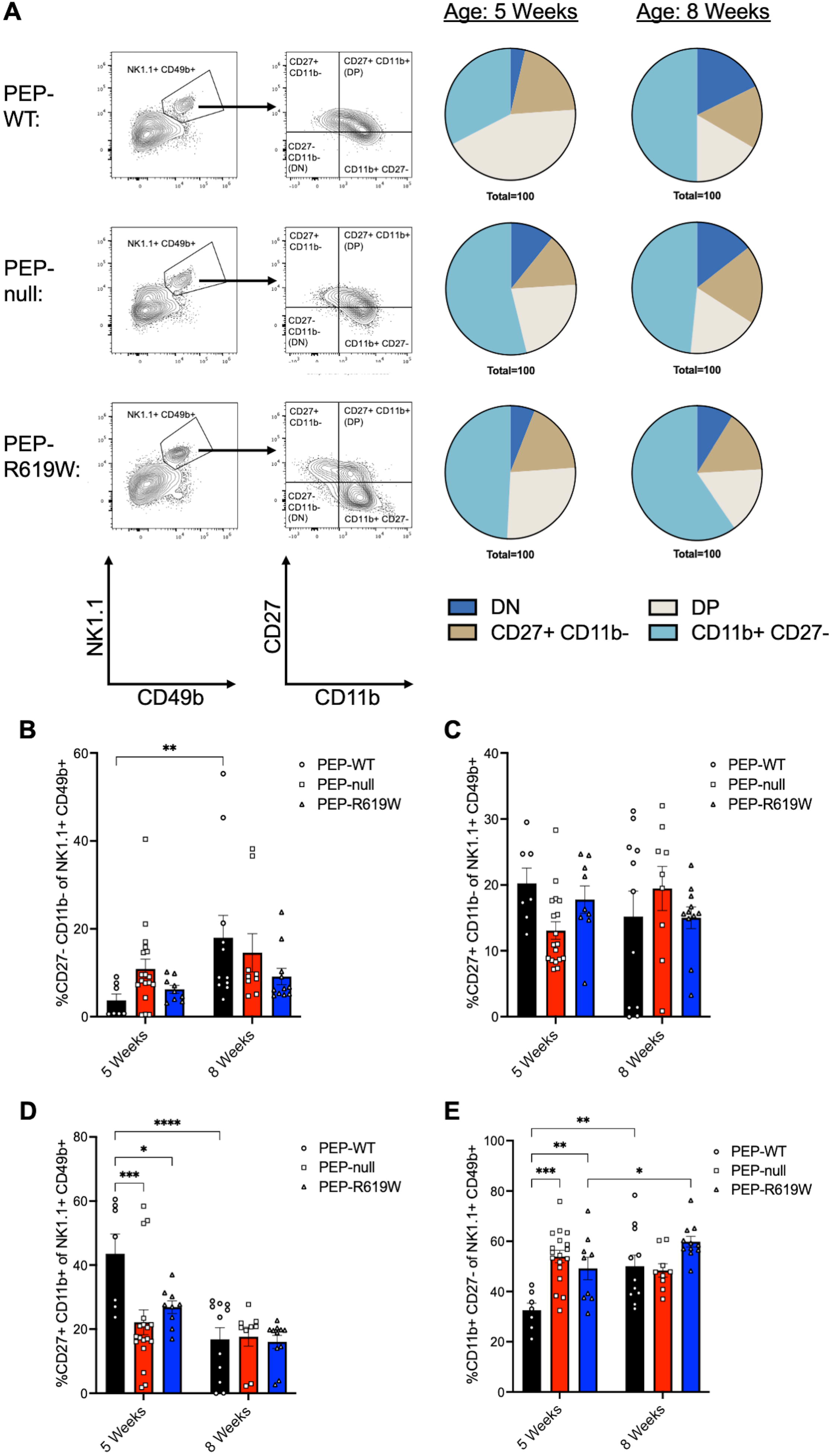
PEP-null and PEP-R619W splenic NK cell repertoires have higher proportions of terminally mature NK cells at 5 weeks of age. Whole spleens were harvested and stained for flow cytometric analysis from 5-week-old or 8-week-old naïve mice as indicated. Representative flow plots from 5-week-old mice are shown from CD3e- CD19- populations, pie charts represent the proportion of the NK1.1+ CD49b+ population that are: CD27- CD11b- (double negative, DN), CD27+ CD11b-, CD27+ CD11b+ (double positive, DP), or CD11b+ CD27-(A). Quantification for frequency (%) of CD27- CD11b- (double negative, DN) (B), CD27+ CD11b- (C), CD27+ CD11b+ (double positive, DP) (D), or CD11b+ CD27- (E) NK cells (NK1.1+ CD49b+). Gating strategy for examined NK cells: Lymphocytes> Single cell x2> autofluorescent-> Live> CD3e- CD19-> NK1.1+ CD49b+> CD27+/- CD11b+/-. Frequencies of flow cytometric data are combined from 5 independent experiments. Each symbol represents an individual mouse. p-values from Two-way ANOVA with Tukey’s Multiple Comparisons are shown for each panel. *p<0.05, **p<0.01, ***p<0.001, ****p<0.0001.

### PEP-R619W NK cells restrict viral replication and enhance IFN**γ** production during coronavirus infection

Given our observation that NK cells are more mature and functional, and there are more of them in PEP-R619W mice during infection, we next wanted to determine if NK cells were necessary for driving disease protection during MHV A59 infection. To test this, we administered an anti-NK1.1 (α-NK1.1) depleting antibody (clone PK136) or anti-IgG isotype control (α-Iso) two days before MHV A59 infection. Subsequent injections were administered every four days after the initial treatment (Fig. 6A). NK depletion was verified at all key time points, both within the spleen and in the peripheral blood (Fig. S6A-E). Loss of NK cells in PEP-R619W mice resulted in a decreased capacity to control virus infection, as viral burden in the liver at 3- and 5-dpi rose to comparable levels of both α-NK1.1 PEP-WT and isotype-treated PEP-WT mice, and were significantly higher than isotype-treated PEP-R619W mice at these time points (Fig. 6B). In addition, isotype-treated PEP-R619W mice maintained the reduced levels of viral titer in the liver compared to isotype-treated PEP-WT mice at these time points (Fig. 6B). These data indicate that PEP-R619W NK cells effectively reduce liver viral load, but PEP-WT NK cells have minimal impact on combating viral replication within the liver. Similar results were detected in the splenic titers (Fig. S6F). These data suggest that in PEP-R619W mice, the increased presence of NK cells in the liver is one of the key innate cells responsible for combating viral replication during the early days of infection. Our data support this conclusion, as all NK-sufficient PEP-R619W mice, including immune-competent (Fig. 1E), Rag1-/- (Fig. 2E), and α-Iso (Fig. 6B) mice, had significantly reduced liver viral titers at days 3 and 5 post-infection compared to PEP-WT counterpart mice. In contrast, this protection was lost in α-NK1.1 PEP-R619W mice (Fig. 6B). Next, we wanted to determine if NK cells were a critical source of IFNγ in our PEP-R619W mice. Upon depletion of NK cells, the levels of IFNγ in liver homogenate at 3 dpi in α-NK1.1 PEP-R619W mice were significantly reduced compared to α-Iso PEP-R619W mice (Fig. 6C). Further, α-Iso PEP-R619W had a significantly higher concentration of IFNγ compared to α-Iso PEP-WT, recapitulating the cytokine profiling data previously shown (Fig. 4G). In conclusion, these data highlight that PEP-R619W enhances NK cell function to protect against MHV A59 viral replication in the liver through, in part, boosting the production of NK cell-derived IFNγ.

**Figure 6:**
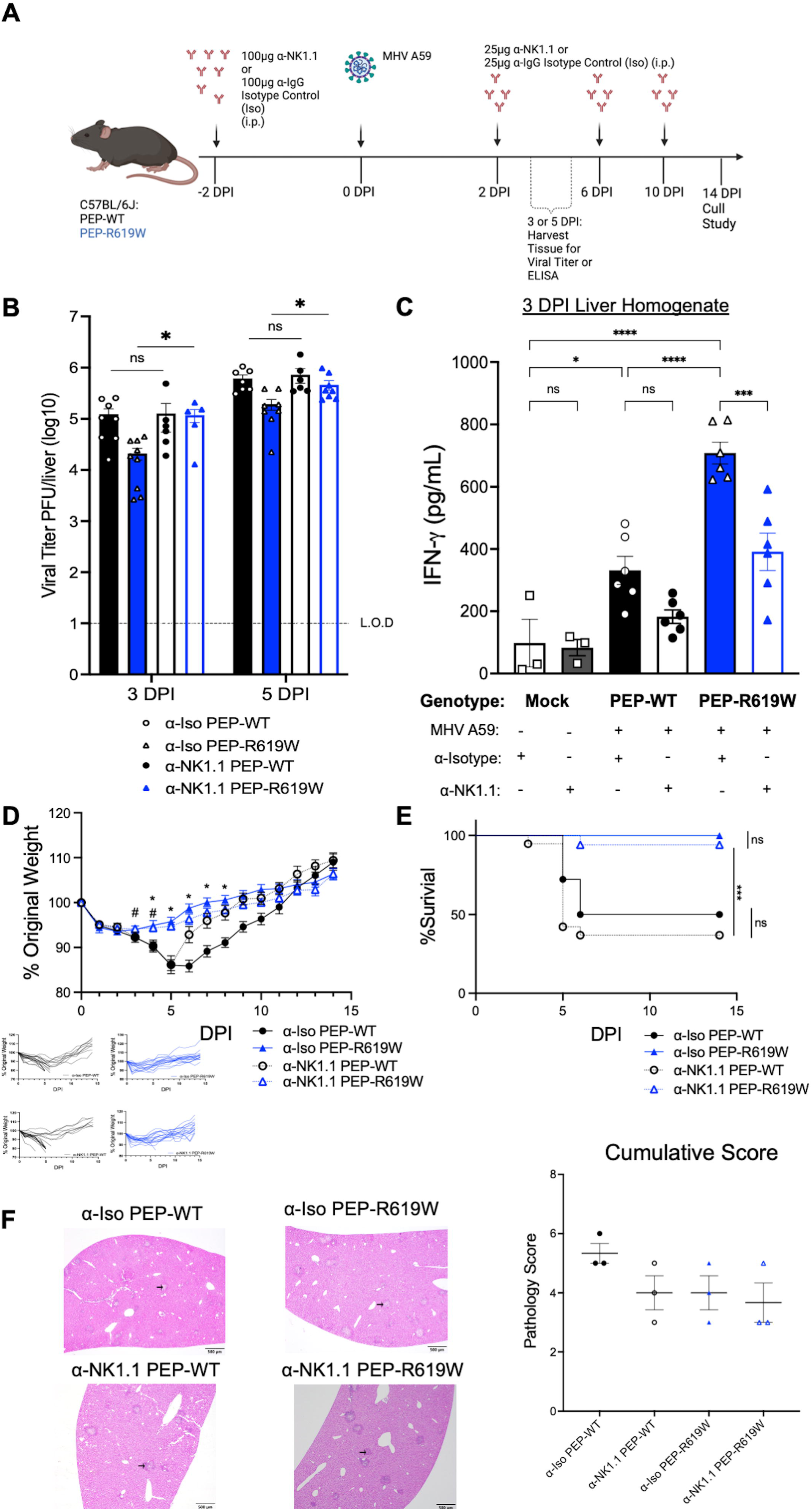
PEP-R619W NK Cells control viral burden and are key producers of IFNγ during MHV A59 infection. PEP-WT and PEP-R619W mice received a natural killer cell depleting antibody (α-NK1.1, Clone PK136) or non-depleting isotype control antibody (α-Iso) as indicated prior- and post-MHV A59 infection (i.p) (A). Following 5e2 PFU MHV A59 infection in α-Iso PEP-WT (black, black circle), α-Iso PEP-R619W (blue, blue triangle), α-NK1.1 PEP-WT (clear, black circle) and α-NK1.1 PEP-R619W (clear, blue triangle) mice, the whole liver was harvested at the indicated dpi, homogenized, and viral titer was quantified via Plaque Assay (B). Mice were injected with PBS (Mock) or infected with 1e5 PFU MHV A59 (i.p), and at 3 dpi, the whole liver was harvested, homogenized, and the levels of IFNγ were determined by ELISA (C). Following 1e5 PFU MHV A59 infection, mice were tracked by weight (D) and survival (E). Respective spider plots for individual mouse weights are shown below (D). Histopathology of the livers of mice at 3 dpi following 1e5 PFU MHV A59 infection, with representative photomicrographs of liver sections exhibiting areas of necrosis, inflammation, and edema/fibrin accumulation (black arrow) and cumulative pathology scores represented as mean + SEM. Scale bar = 500 μm (F). Each symbol represents an individual mouse for whole liver titer (B) and IFNγ ELISA (C). Viral titers are pooled data from 2 independent experiments at each time point, comparisons are shown from multiple unpaired T-Tests, α-Iso PEP-WT vs α-NK1.1 PEP-WT, α-Iso PEP-R619W vs α-NK1.1 PEP-R619W, *p<0.01. The dashed line indicates the limit of detection (L.O.D.). IFNγ ELISA is representative of two independent experiments, p-values from One-way ANOVA with Tukey’s Multiple Comparisons are shown, *p<0.05, **p<0.01, ***p<0.001, ****p<0.0001. Mock mice are a mixture of both genotypes. Weight loss and survival studies were pooled from 2 individual experiments. Group sizes were α-Iso PEP-WT n = 18, α-Iso PEP-R619W n = 17, α-NK1.1 PEP-WT n = 19, and α-NK1.1 PEP-R619W n = 17. For weight loss statistical analysis (D), significant p-values from One-way ANOVA with Tukey’s Multiple Comparisons at each dpi are represented by # comparing α-NK1.1 PEP-WT vs α-NK1.1 PEP-R619W, or * comparing α-Iso PEP-WT vs. α-Iso PEP-R619W. Log-rank Mantel-Cox Test (survival) (E), ***p<0.001. For histopathological analysis, each dot represents an individual mouse, with p-values from One-way ANOVA with Tukey’s Multiple Comparisons shown, *p<0.05.

### PEP-R619W NK cells do not mediate weight loss, survival, or liver histopathology during coronavirus infection

Since PEP-R619W NK cells mediate viral titer at early days post-infection and are a key source of IFNγ, we next wanted to determine if they were necessary for mediating weight loss and survival during MHV A59 infection. Following 1e5 PFU MHV A59 infection, NK cell depletion in either PEP-WT or PEP-R619W mice did not result in changes in weight loss, survival, or in liver histopathology compared to NK-sufficient PEP-WT and PEP-R619W counterparts (Fig. 6D-F and Fig. S6G).

Overall, these data indicate that NK cells in PEP-R619W mice play a specific role in combating MHV A59 infection by restricting viral replication and are critical producers of IFNγ within the primary site of replication, the liver. However, these NK cells are dispensable for protection against weight loss, survival, and liver damage. While NK cell depletion in PEP-R619W mice increased viral load and reduced liver IFNγ concentration, the same depletion in PEP-WT mice did not affect liver viral titer or IFNγ production. This disparity can potentially be attributed to the reduced number and impaired function of M-cNK cells in the livers of PEP-WT mice at 3 dpi (Fig. 3D-G, Fig. 4A and 4B), making their absence less impactful on viral replication.

## DISCUSSION

Despite the strong genetic link between the minor allele of *PTPN22* and the development of various autoimmune diseases, this allele persists in the human population. A potential theory lies in the fact that the mechanisms driving heightened immune responses, while pathogenic in the context of autoimmunity, are likely protective when defending against infections and cancer. Previous studies support this reasoning, as PEP-R619W mice exhibit enhanced antiviral immunity during chronic LCMV-cl13 infection and increased antitumor immunity, resulting in reduced disease severity compared to PEP-WT mice (18, 19). Similar results in PEP-null mice have been reported during LCMV-cl13 infection (30–32). However, these diseases are chronic. Thus, we were interested in determining if PEP-null and/or PEP-R619W mice would also have improved disease outcomes during acute, moribund coronavirus infection, MHV A59.

Our results show that PEP-R619W and PEP-null do not always phenocopy each other. Regardless of whether PEP-null mice were fully immune-competent or on a Rag1-/- genetic background, their phenotypes were comparable to PEP-R619W mice after 5e4 PFU infection, but demonstrated increased susceptibility following 1e5 PFU infection and mirrored the outcomes observed in PEP-WT mice (Figs. 1 and 2). Despite similar splenic innate profiles and NK activation between PEP-null and PEP-R619W mice at 3 dpi following 5e4 PFU infection, after 1e5 PFU MHV A59 infection, PEP-null splenic NK cell phenotypes were comparable to PEP-WT (Fig. S3 and S5). These findings add to the current literature, highlighting that there are differences between PEP-null and PEP-R619W. While PEP-null and PEP-R619W mice may be similar in certain scenarios, such as in LCMV-cl13 antiviral immunity, there are other examples where they can differ (19–21, 27, 30–32). This may lie in cellular and/or molecular differences between these strains. For example, previous studies show that PEP-null mice have more Tregs than PEP-WT and PEP-R619W mice, and that the PEP-R619W mutation does not directly impact phosphatase activity (3, 4, 9, 23, 24, 26, 27, 29). Instead, the mutation disrupts molecular interactions with other proteins, such as TRAF3 and Csk, leading to altered immune function (3, 4, 9, 23, 24, 26, 27, 29). The precise ways in which established molecular interactions and other potential mechanisms that influence innate immune cell function diverge between PEP-null and PEP-R619W mice are poorly understood. Further investigation into these mechanisms would provide crucial insight into the varying outcomes observed between these strains during MHV A59 infection.

Our data shows that PEP-R619W protects against murine coronavirus, MHV A59, associated disease (Fig. 1). This is paired with reduced liver viral titer at early days post-infection, and accelerated clearance compared to PEP-WT mice (Fig. 1). This protection in PEP-R619W mice was partially mediated by an enhanced innate immune response. This is further strengthened by the results of our studies using Rag1-/- models, as Rag1-/- PEP-R619W mice had reduced disease compared to Rag1-/- PEP-WT mice (Fig. 2). These data indicate that PEP-R619W differentially modulates the innate immune response to defend against acute MHV A59 infection (Figs. 1 and 2). These data are further supported by the immune phenotyping of both the spleen and liver at 3 dpi, where PEP-R619W mice had more innate immune cells compared to PEP-WT mice (Fig. 3, Fig. S3 and S4).

We interrogated the immune response in the liver following 1e5 PFU infection to define how PEP-R619W uniquely enhanced innate antiviral immunity. Notably, PEP-R619W mice had more NK cells, DCs, and macrophages compared to PEP-WT and PEP-null mice (Fig. 3D). However, we detected no differences in type-I interferon (IFN-I) production, which is necessary for survival following MHV A59 infection (54). There were no changes in the pDC population, which are critical producers of IFN-I (Fig. 3D), or in the systemic and liver homogenate levels of IFNα and IFNβ (Fig. 4C, E, F, H, and I) between infected genotypes, indicating that PEP-R619W protects through alternative mechanisms, such as enhancing NK cell function. NK cells can directly lyse infected cells through the release of perforin and granzyme B, are the primary producers of IFNγ during the innate immune response, and these functions are enhanced as NK cells mature (46, 47, 55, 56). These effector functions are necessary for combating murine cytomegalovirus (MCMV) disease and viral replication and have been specifically shown to be vital in driving the innate immune response in the liver post-infection (57–61). As such, we more closely interrogated the NK cell compartment in PEP-WT and PEP-R619W mice post-MHV A59 infection.

We investigated the role of NK cells during MHV A59 infection and discovered that: 1) there were more mature conventional NK (M-cNK) cells in the spleen and liver of PEP-R619W mice (Fig. 3D-G and Fig. S3D, S3E), 2) PEP-R619W enhanced M-cNK effector function and polyfunctionality compared to PEP-WT (Fig. 4A and 4B), and 3) PEP-R619W NK cells, but not PEP-WT NK cells, are necessary for restricting viral replication in the liver at 3 and 5 dpi (Fig. 6B) and are a critical source of IFNγ (Fig. 6C). Taken together, these results indicate that PEP-R619W augments M-cNK cell antiviral function in the liver during MHV A59 infection, leading to reduced viral replication, but PEP-R619W M-cNK cells do not directly mediate weight loss, survival, or liver pathology (Fig. 6D-F). These findings add to recent work and highlight the protective role of NK cells at the primary site of coronavirus infection, as NK cells are critical for combating SARS-CoV-2 replication in both human and non-human primate lungs (62–64). However, our data also demonstrates that viral burden and disease symptoms are not always directly correlated. This has been observed in previous MHV A59 studies, as well as during chronic LCMV-cl13 infection, where weight loss is driven by CD8 T-cell immunopathology (34, 65).

We are one of the first groups to investigate the function of PEP in NK cells. The molecular role of PEP in NK cells remains undefined, despite being the immune cells with the highest Lyp/PEP expression in humans and mice (66–68). Our results show that at 5 weeks of age, PEP-R619W mice have more mature NK cells post-MHV A59 infection at 3 dpi in the spleen and liver compared to PEP-WT (Fig. S3 and Fig. 3). This phenotype persists at homeostasis, where both PEP-null and PEP-R619W mice had a higher proportion of terminally mature (CD11b+ CD27-) NK cells in the spleen compared to PEP-WT counterparts (Fig. 5). Despite these findings, the molecular functions of PEP within NK cells, and how the PEP-R619W variant accelerates maturity and enhances effector functions during infection, remains unknown. These results warrant additional studies defining the molecular mechanisms by which PEP and PEP-R619W regulate NK cell maturity and function at homeostasis and during disease.

In conclusion, our results indicate that the autoimmunity-associated allele of *PTPN22* enhances innate antiviral immunity against acute murine coronavirus infection, MHV A59, and adds to the growing literature supporting the protective role of this allele during viral and tumorigenic challenges (18, 19). We have shown a divergence in protection against MHV A59 infection in PEP-null mice in an inoculation dose-dependent manner, whereas PEP-R619W was protective regardless of infection dose. Additionally, we showed that PEP-R619W boosts innate cell infiltration, enhances NK cell maturation and function, and, overall, reduces disease following murine coronavirus infection. These data highlight that PEP-R619W uniquely modifies immune function through distinct mechanisms compared to both PEP-WT and PEP-null, especially in NK cells. These findings provide further rationale for testing the protective capacity of PEP-R619W in additional infection models and defining how PEP-R619W augments immunity in a context-dependent manner.

## MATERIALS AND METHODS

### Ethics Statement

The University of Kansas (KU) Institutional Animal Care and Use Committee (IACUC) reviewed and approved all animal studies under protocol number AUS 278-01.

### Mice

Both male and female mice at 5 weeks of age were used in all studies unless otherwise noted. We do not observe sex-based differences in these studies. C57BL/6J mice and C57BL/6 Rag1-/- mice were originally purchased from Jackson Labs and then bred and maintained in the University of Kansas Animal Care Unit (ACU) facilities. *Ptpn22* knockout (PEP-null) and *Ptpn22* R619W (PEP-R619W) mice were generated using CRISPR/Cas9 technology using methods previously reported (15, 18, 19, 69). PEP-null Rag1-/- and PEP-R619W Rag1-/- mice were derived as previously described (18). IFNγ KO (RRID: IMSR_JAX:002287) and IFNγR KO (RRID: IMSR_JAX:003288) mice were purchased from Jackson Labs and were housed and maintained in the University of Kansas ACU facilities as detailed above.

### Cell Culture

#### HeLa-MHVR and 17Cl-1 Cells

HeLa cells expressing the MHV receptor CEACAM1a (HeLa-MHVR) were cultured in Advanced Dulbecco’s modified Eagle’s medium (DMEM) supplemented with 10% FBS, 1% L-glutamine, and 1% Penicillin/Streptomycin, and 17Cl-1 cells were grown in DMEM supplemented with 5% FBS, 5% Tryptose Phosphate Broth, and 2% Penicillin/Streptomycin (70) (71).

#### Bone Marrow Dendritic Cells (BMDC) and Macrophages (BMM)

To generate BMDCs, bone marrow cells were isolated from PEP-WT, PEP-null, and PEP-R619W mouse femurs. Following isolation, cells were cultured in Advanced DMEM supplemented with 10% FBS, 1% L-glutamine, 1% Penicillin/Streptomycin, and 100ng/mL FLT3L (Stem Cell Technologies, Vancouver, CA). For BMMs, bone marrow cells were isolated and cultured in Roswell Park Memorial Institute medium (RPMI) supplemented with 10% FBS, 1% L-glutamine, and 1% Penicillin/Streptomycin with 50ng/mL M-CSF (Stem Cell Technologies, Vancouver, CA) for 8 days. An additional 10 mL of M-CSF-containing media was added to BMM cultures on day 3, and on day 8, differentiated cells were harvested, counted, and replated for further assays (72).

### MHV A59 Production, Titer, and Infections

MHV A59 stocks were grown in 17Cl-1 cells, and viral titers were determined on HeLa cells expressing the MHV receptor CEACAM1a (HeLa-MHVR), as previously described (42, 70, 71, 73, 74). Briefly, HeLa-MHVRs were seeded in 12-well plates the day before infection. The following day, once the HeLa-MHVRs were grown to 90-95% confluency, 10-fold dilutions of titer samples were prepared, and the HeLa-MHVRs were inoculated for a 45-to-60-minute adsorption phase, followed by 1 mL overlay per well of 1:1 mixture of 1.2% Agarose and 2x DMEM (10% FBS, 1% L-Gln, 1% Pen/Strep). Once solidified, 500 µL D2 media (DMEM, 2% FBS, 1% L-Gln, 1% Pen/Strep) was added to the overlay. The plaques were counted and recorded 48 hours later for viral titer (PFU/sample) calculation.

For *in vivo* infections, mice were inoculated with 5e2 PFU, 5e4 PFU, or 1e5 PFU MHV A59 in a final volume of 200 µL sterile PBS via intraperitoneal (i.p.) injection (PBS alone for “mock” mice). Throughout all infections, mice were weighed daily and euthanized if weight loss was greater than 20% of the original starting weight. At peak infection (days 5-8 post-infection), mice were checked twice daily to provide more frequent monitoring. To determine viral load, the whole liver and spleen were harvested and homogenized in 5 mL or 1 mL sterile DMEM, respectively, centrifuged to remove cellular debris, and stored at -80^0^C. The viral titer was determined on HeLa-MHVR cells as detailed above.

For *ex vivo* Bone Marrow Dendritic Cell (BMDC) and Bone Marrow Macrophage (BMM) infections, cells were inoculated at a multiplicity of infection (MOI) of 0.1 with a 45-to-60-minute adsorption phase. The viral inoculum was aspirated, and 1 mL of advanced DMEM supplemented with 10% FBS, 1% L-glutamine, 1% Penicillin/Streptomycin or RPMI supplemented with 10% FBS, 1% L-glutamine, and 1% Penicillin/Streptomycin, for BMDC or BMM, respectively, was added per well and incubated for 24 hours. At 24 hours post-infection, the cells were scraped and resuspended before being placed in the -80^0^C freezer overnight. Following freeze-thaw, the virus from the supernatant and cells was combined and tittered on HeLa-MHVRs as described above.

### Leukocyte Isolation and Flow Cytometry

Mice were euthanized at indicated time points in line with AVMA and KU IACUC guidelines by CO2 inhalation followed by cervical dislocation before tissue harvest.

The spleen was excised at indicated days post-infection, smashed over 40 µm filters, rinsed with DMEM, and centrifuged to remove debris. The cell pellet was resuspended in 2 mL ACK red cell lysis buffer, incubated at RT for 2 minutes, quenched with 10 mL DMEM, and centrifuged. The resultant pellet was resuspended in 2 mL advanced DMEM supplemented with 10% FBS, 1% L-glutamine, 1% Penicillin/Streptomycin to generate a single cell suspension.

Whole livers were harvested following cardiac perfusion with 15-20 mL sterile PBS and placed in 3 mL DMEM supplemented with 10% FBS on ice. Livers were digested and processed using a murine liver dissociation kit (Miltenyi Biotec, Bergisch Gladbach, North Rhine-Westphalia, Germany) per the manufacturer’s protocol. Following centrifugation, the cell pellet was resuspended in 1 mL sterile DMEM (1% L-glutamine), gently overlaid onto a 20 mL 33% percoll gradient in PBS, and centrifuged at 800xg for 25 minutes at room temperature (RT) (75). After aspiration, the cell pellet was resuspended in 2 mL Advanced DMEM (supplemented with 10% FBS, 1% L-glutamine, 1% Penicillin/Streptomycin) to generate a single cell suspension.

To harvest peripheral blood for NK Depletion verification, 50 µL of blood was acquired via retro-orbital (r.o.) bleeds using heparinized capillary tubes. 10 µL of blood was plated in a 96-well plate for each sample. 100 µL of ACK red cell lysis buffer was added and incubated at RT for 2 minutes, quenched with 100 µL Advanced DMEM (supplemented with 10% FBS, 1% L-glutamine, 1% Penicillin/Streptomycin), centrifuged, and resuspended in 200 µL of media to generate a single cell suspension.

All single cell suspensions were counted using Vi-CELL BLU (Beckman Coulter, Brea, CA). The concentration of each sample was adjusted to plate 2e6 cells/well in round-bottom 96-well plates for antibody staining and flow cytometric analysis.

All flow cytometry was performed on the Cytek Aurora 5 laser system (355 nm [UV], 405 nm [Vio], 488 nm [Blue], 561 nm [YG], and 640 nm [Red]) spectral flow cytometer (Fremont, CA, US). Single stain control Biolegend Compensation Beads were used for unmixing (San Diego, California, U.S). Unmixed files were analyzed using FlowJo Software (BD Biosciences, San Diego, California). Antibodies used in varying combinations (depending upon the experiment) are shown below in the Supplementary Materials (Table S1).

All samples were incubated with diluted Fc block (Biolegend) for 10 minutes and then stained for 30 minutes at 4^0^C in the dark for surface marker staining in serum-free PBS. For intracellular cytokine staining, cells were incubated with brefeldin A (BFA, Thermo Scientific) for four hours at 37^0^C before surface staining. BD Cytofix and Permwash kit or Tonbo Foxp3 Fix/Perm kit were used according to manufacturers’ protocols for intracellular cytokine or intranuclear transcription factor staining, respectively.

### Cytokine and Chemokine Quantification

500 µL of blood was acquired via retro-orbital (r.o.) bleeds using non-heparinized capillary tubes before euthanasia. After incubating at RT for 30 minutes, the blood was centrifuged at 13,000 RPM for 10 minutes at RT, and the serum supernatant was collected and stored at -80^0^C for future analysis. As described above, the whole liver was homogenized in 5 mL DMEM, centrifuged to remove cellular debris, and stored at -80^0^C for future analysis. Serum and liver homogenate antiviral cytokines and chemokines were quantified using a LEGENDplex assay (Biolegend), with the serum being diluted 1:2 as described in the manufacturer protocol. In a separate experiment, a portion of the left lateral liver lobe was harvested, weighed, homogenized, and the concentration of IFNβ was determined using the Mouse IFN-Beta ELISA Kit (PBL Assay Science). The levels of IFNγ were determined using LEGEND MAX™ Mouse IFNγ ELISA Kit (Biolegend) in NK cell depletion experiments following whole liver harvest and homogenization at indicated time points.

### Natural Killer (NK) cell depletion

Depletion of NK cells was achieved through i.p. injection of 100 µg anti-NK1.1 monoclonal antibodies (clone PK136) two days before infection and followed by 25 µg anti-NK1.1 injections every four days to sustain depletion (Bio-X-Cell). In parallel, separate mice were administered control IgG injections at the same dosage and timings as detailed above (Bio-X-Cell). The experimental outline is shown in Supplementary Figure 6.

### Histopathology

Whole livers were harvested following cardiac perfusion with 15-20 mL sterile PBS, followed by 10-15 mL of 4% PFA diluted in PBS, and placed in 20 mL of 10% PFA diluted in PBS (minimum ratio of 1 gram tissue:10 mL 10% PFA). After 72 hours at 4^0^C in the dark, a representative section from each lateral and medial (right and left) lobe was cut and processed for routine hematoxylin and eosin (H&E) staining. The liver lesions were blindly scored by an American College of Veterinary Pathology Board-certified pathologist. The lesions scored were inflammation, necrosis, and edema/fibrin on a scale of 0–10% (score 1), 10–40% (score 2), 40– 70% (score 3), and >70% (score 4). Cumulative scores were obtained for each mouse and presented as total pathology score.

### Statistical Analysis and Graphing

All graphs and appropriate statistical analysis for data were done in GraphPad Prism (La Jolla, CA). The type of statistical tests used are listed in each figure legend. Data was considered statistically significant if the p-value<0.05. Data with a p-value>0.05 was determined non-significant, and pairwise comparisons were either not labeled or represented with “ns.” Figure legends indicate if data are from pooled experiments or are representative of multiple studies. Most graphs display mean + SEM, unless otherwise stated.

## Supplementary Materials

Figure S1: MHV A59 disease is attenuated in 8-week-old mice, and PEP-R619W does not impact splenic titer or restrict viral replication in myeloid cells.

Figure S2: Splenic titers are unchanged in Rag1-/- mice regardless of PEP genotype.

Figure S3: PEP-R619W increases the splenic innate immune population.

Figure S4: Representative gating strategy for different liver immune cell populations and additional quantification of immune cells.

Figure S5: Infection dose-dependent impact on splenic mature NK function.

Figure S6: NK depletion protocol and NK-depleted spleen titer and histopathology scores.

Table S1: Flow Cytometry Antibodies

## ACKNOWLEDGMENTS

We thank the KU Flow Cytometry Core, KU Animal Care Unit, and all members of the Orozco lab for their discussions and assistance. We also thank Dr. Hans Dalton for his edits on this manuscript.

## Funding Sources

NIH T32 Chemical Biology T32 Training Grant GM13206 (AB)

KU Chemical Biology of Infectious Disease (COBRE) National Institute of General Medical Sciences (NIGMS) of the National Institutes of Health under award number P20GM113117 (RCO, ARF)

KU Center for Genomics Seed Grant (RCO)

Funds from the University of Kansas (RCO)

NIGMS R35GM138029 (ARF)

NIH T34 GM062232 (KJLHR)

## Author contributions

Conceptualization: AMB, ARF, RCO

Methodology: AMB, KJLHR, TC, CMK, SM, ARF, RCO

Investigation: AMB, KJLHR, NS, CMK, SM, RCO

Visualization: AMB, KJLHR, SM, ARF, RCO

Funding acquisition: AMB, KJLHR, RCO

Project administration: RCO Supervision: SM, ARF, RCO

Writing – original draft: AMB, SM, ARF, RCO

Writing – review & editing: AMB, KJLHR, NS, TRC, CMK, SM, ARF, RCO

## Competing interests

Authors declare that they have no competing interests.

## Data and materials availability

All data are available in the main text or the supplementary materials. Raw data is available upon request.

## Notes

### Competing Interest Statement

The authors have declared no competing interest.

### Summary of Updates

This file has been updated to include some new data (SF1) and has revised clarity for reading and interpretation of results.

