## Supplemental Figure for "An Autoimmunity-associated allele of *PTPN22* enhances innate antiviral immunity to protect against acute coronavirus infection"

**Supplementary Materials**

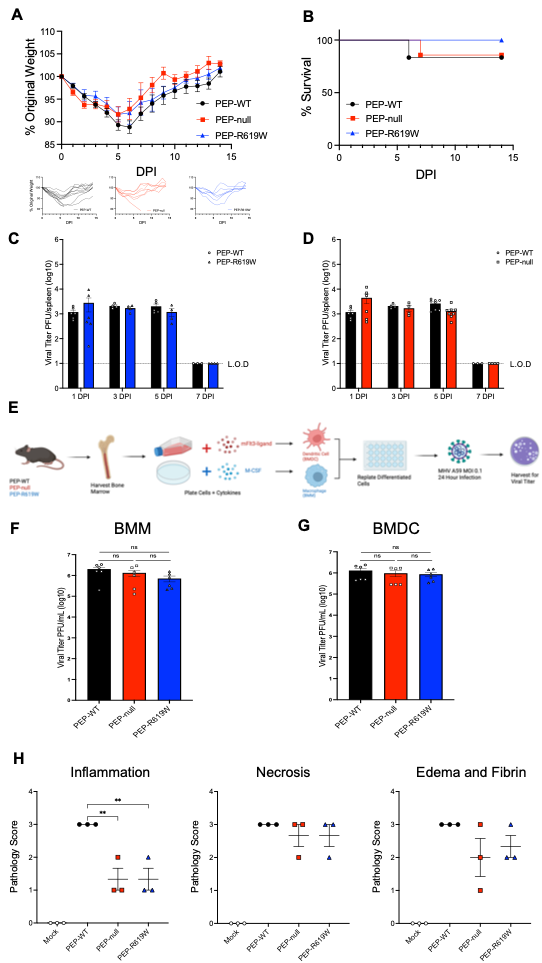
**Fig. S1**

**Figure S1: MHV A59 disease is attenuated in 8-week-old mice, and PEP-R619W does not impact splenic titer or restrict viral replication in myeloid cells.**

8-week-old C57BL/6J mice with *Ptpn22* wild type (PEP-WT, black circle), *Ptpn22* knockout (PEP-null, red square), or the *Ptpn22* autoimmune-associated allele (PEP-R619W, blue triangle), were infected with 1e5 PFU (A, B) MHV A59 (i.p). Mice are tracked by weight (A) and survival (B). Respective spider plots for individual mouse weights are shown below (A). To measure viral burden, spleen homogenates from infected (5e2 PFU MHV A59) PEP-WT, PEP-null, and PEP-R619W mice were used for plaque assay (C, D). Experimental schematic for deriving bone marrow-derived macrophages (BMMs) and bone marrow-derived dendritic cells (BMDCs) (E). The viral titer was quantified via Plaque Assay for BMMs (F) and BMDCs (G). Liver histopathology scores of individual parameters (inflammation, necrosis, and edema/fibrin) at 3 dpi following PBS injection (Mock) or 1e5 PFU MHV A59 infection are shown (H). For weight loss and survival studies, group sizes were PEP-WT n = 12, PEP-null n = 7, and PEP-R619W n = 6. For whole spleen titer (C, D), each symbol represents an individual mouse. The dashed line indicates the limit of detection (L.O.D.). Certain PEP-WT mice are represented in both graphs, as some experiments had all three genotypes, whereas others contained only PEP-WT vs. PEP-null or PEP-R619W. Combined data from 2 independent experiments, multiple unpaired T-Tests, PEP-WT vs PEP-R619W, PEP-WT vs PEP-null, at each time point. For BMM and BMDC titer, each MHV A59 infection was performed at MOI = 0.1. Data are shown from two independent experiments with 3 technical replicates per experiment. For histopathological analysis, mock mice received sterile PBS injection only, and are a mixture of all three genotypes. Each dot represents an individual mouse, with p-values from One-way ANOVA with Tukey’s Multiple Comparisons shown, *p<0.05, **p<0.01.

**Fig. S2**

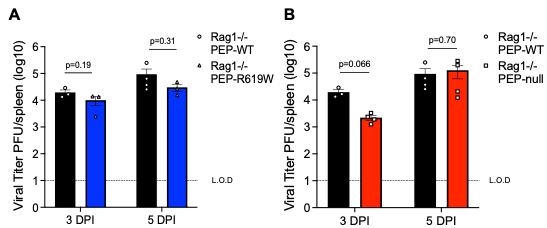
**Figure S2: Splenic titers are unchanged in Rag1-/- mice regardless of PEP genotype.**

To measure viral burden, liver homogenates from infected (5e2 PFU) Rag1-/- PEP-WT (black), Rag1-/- PEP-null (red), and Rag1-/- PEP-R619W (blue) mice were used for plaque assay (A, B). Each symbol represents an individual mouse. The dashed line indicates the limit of detection (L.O.D.). Certain Rag1-/- PEP-WT mice are represented in both graphs, as some experiments had all three genotypes, whereas others contained only Rag1-/- PEP-WT vs Rag1-/- PEP-null or Rag1-/- PEP-R619W. Multiple unpaired T-Tests, PEP-WT vs PEP-R619W, PEP-WT vs PEP-null, at each time point.

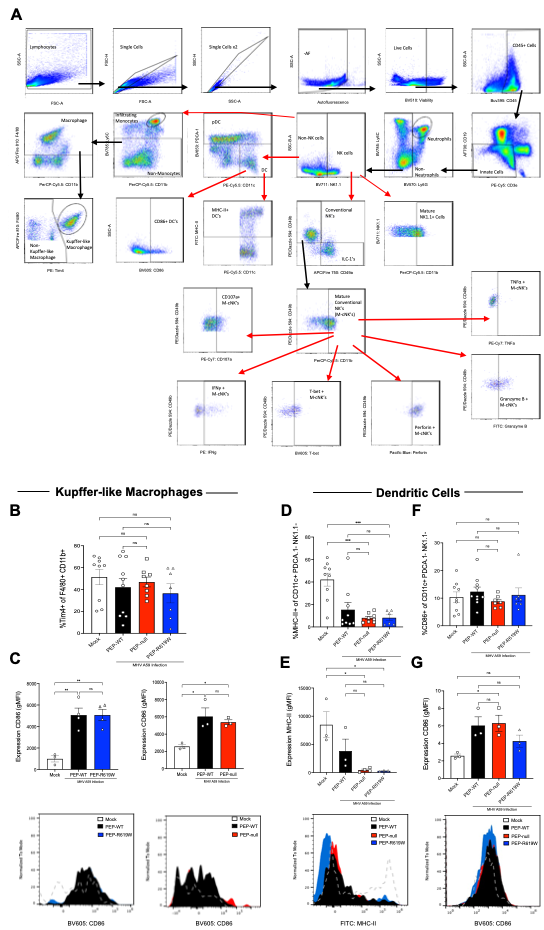
**Fig. S3**

**Figure S3: PEP-R619W increases the splenic innate immune population.**

Representative gating strategies to identify splenic immune cell populations of interest (A). Populations are labeled on flow cytometry plots, where black arrows indicate the down-gating of the population from which they are drawn from, and red arrows indicate the down-gating from one population to multiple populations. Following PBS injection (Mock, white bar, circle) or 5e4 PFU MHV A59 infection (i.p) in PEP-WT (black bar, circle), PEP-null (red bar, square), PEP-R619W (blue bar, triangle) mice, at 3 dpi the spleen was harvested and antibody stained for flow cytometric analysis. The absolute count of splenocytes for all genotypes (B). Frequencies (%) of CD3e- CD19- of Live cells (C). Frequency (%) of several innate cells of Live cells is shown (D). The gating strategy for all innate subsets is: Lymphocytes> Single cell x2> autofluorescent-> Live> CD3e- CD19-> Ly6G- Ly6C+/- (non-neutrophils). NK Cells: innate subsets> NK1.1+. DCs: innate subsets> CD11c+ PDCA-1-. pDCs: innate subsets> PDCA-1+ CD11c+/-. Monocytes: innate subsets> Ly6Chi CD11b+. Macrophage: innate subsets> Ly6C- CD11b+/-> CD11b+> F4/80+. Neutrophils: Lymphocytes> Single cell x2> autofluorescent-> Live> CD3e- CD19-> Ly6G+ Ly6C+. Frequency (%) of CD11b+ (mature) NK cells (E). Frequency (%) Splenic CD8α+ cDCs (F) and expression (gMFI) of CD80 (G) and CD86 (H) with representative histograms to the right. Frequency (%) CD8α- cDCs (I) and expression (gMFI) of CD80 (J) and CD86 (K) with representative histograms to the right. Quantification of flow cytometric data was pooled from 3 independent experiments. Each symbol represents an individual mouse. Mock mice are a mixture of all 3 genotypes. p-values from One-way ANOVA with Tukey’s Multiple Comparisons are shown for each panel. *p<0.05, **p<0.01, ***p<0.001, ****p<0.0001.

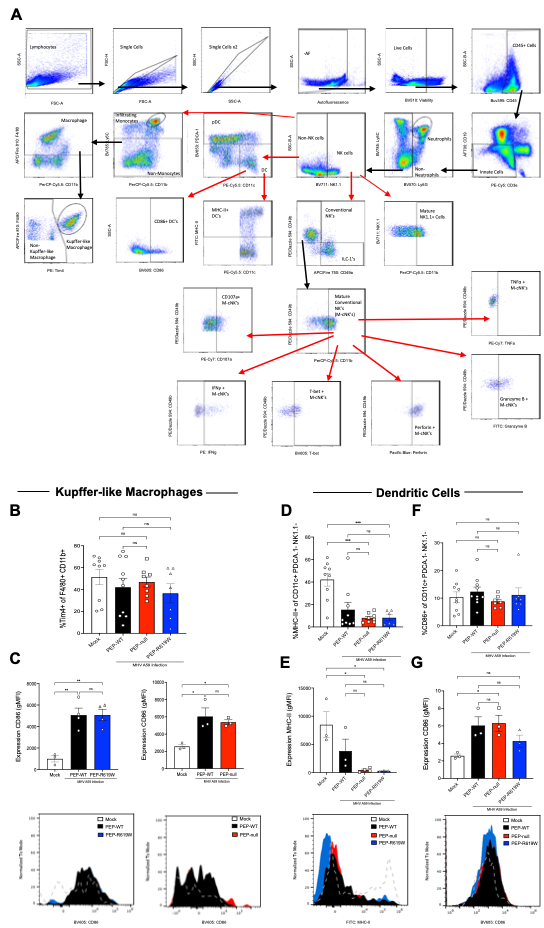
**Fig. S4**

**Figure S4: Representative gating strategy for different liver immune cell populations and additional quantification of immune cells.**

Representative gating strategies to identify liver immune cell populations of interest presented in Figure 3 (A). Populations are labeled on flow cytometry plots, where black arrows indicate the down-gating of the population from which they are drawn from, and red arrows indicate the down-gating from one population to multiple populations. Following PBS injection (Mock, white bar, circle) or 1e5 PFU MHV A59 infection (i.p) in PEP-WT (black bar, circle), PEP-null (red bar, square), PEP-R619W (blue bar, triangle) mice, at 3 dpi the liver was perfused, harvested, and antibody stained for flow cytometric analysis. Tim4+ Macrophages (Kupffer-like cells) frequency (%) (B) and expression of CD86 (gMFI) in Tim4+ Macrophages (C), with representative histograms below. The first plot (left) comparing Mock, PEP-WT, and PEP-R619W and the second plot (right) comparing Mock, PEP-WT, and PEP-null are representative of two separate independent experiments. The gating strategy for Kupffer-like (Tim4+ macrophages) is: Lymphocytes> Single cell x2> autofluorescent-> Live> CD45+> CD3e- CD19-> Ly6G- Ly6C+/- (non-neutrophils)> NK1.1-> Ly6C- CD11b+/-> F4/80+ CD11b+> Tim4+. MHC-II+ DCs frequency (%) (D) and expression of MHC-II (gMFI) in DCs (E) with a representative histogram below. CD86+ DCs frequency (%) (F) and expression of CD86 (gMFI) in DCs (G) with a representative histogram below. The gating strategy for MHC-II+ DCs or CD86+ DCs is: Lymphocytes> Single cell x2> autofluorescent-> Live> CD45+> CD3e- CD19-> Ly6G- Ly6C+/- (non-neutrophils)> NK1.1-> CD11c+ PDCA-1-> MHC-II+ or CD86+. Quantification of flow cytometric data was pooled from 2 independent experiments. Each symbol represents an individual mouse. Mock mice are a mixture of all 3 genotypes. p-values from One-way ANOVA with Tukey’s Multiple Comparisons are shown for each panel. *p<0.05, **p<0.01, ***p<0.001, ****p<0.0001.

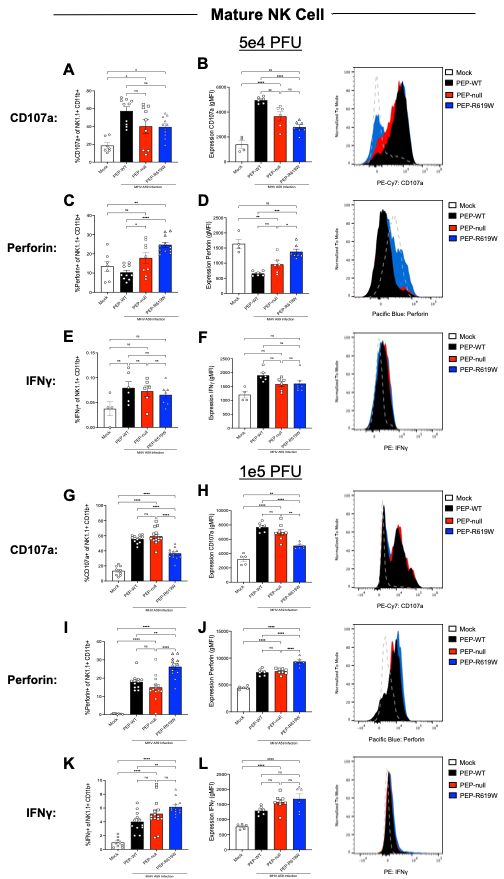
**Fig. S5**

**Figure S5: Infection dose-dependent impact on splenic mature NK function.**

Following PBS injection (Mock, white bar, circle) or 5e4 PFU (A-F) or 1e5 PFU (G-L) MHV A59 infection (i.p) in PEP-WT (black bar, circle), PEP-null (red bar, square), and PEP-R619W (blue bar, triangle) mice, at 3 dpi the spleen was harvested, surface and intracellularly antibody stained for flow cytometric analysis. Frequency (%), expression (gMFI), and representative histogram to the right of CD107a (A-B, G-H), Perforin (C-D, I-J), and IFNγ (E-F, K-L) in Mature NK cells, for 5e4 PFU and 1e5 PFU MHV A59 infections, respectively. The gating strategy for Mature NK cells and effector markers was: Lymphocytes> Single cell x2> autofluorescent-> Live> CD3e- CD19-> Ly6G- Ly6C+/- (non-neutrophils)> NK1.1+> CD11b+> CD107a+ or Perforin+ or IFNγ+. Frequencies of flow cytometric data are combined from 2 independent experiments, gMFI is representative images from 2 independent experiments at each dose. Each symbol represents an individual mouse. Mock mice are a mixture of all 3 genotypes. p-values from One-way ANOVA with Tukey’s Multiple Comparisons are shown for each panel. *p<0.05, **p<0.01, ***p<0.001, ****p<0.0001.

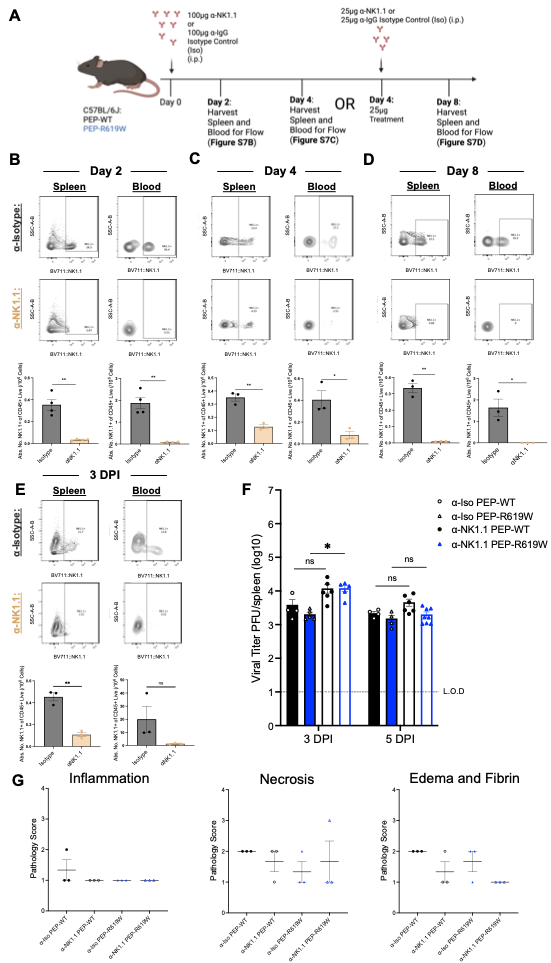
**Fig. S6**

**Figure S6: NK depletion protocol and NK-depleted spleen titer and histopathology scores.**

Experimental timeline detailing the concentration of α-NK1.1 antibody (Clone PK136) and frequency of doses (A). NK cell depletion representative flow plots and absolute counts of NK1.1+ cells (per 10^6^ cells) are shown for the spleen and peripheral blood from naïve mice at day 2 (B) and day 4 (C) post-100μg α-NK1.1 or α-Iso injection (i.p). Alternatively, naïve mice were administered 100μg α-NK1.1 or α-Iso injection (i.p), and on day 4, an additional dose of 25μg α-NK1.1 or α-Iso injection was given. NK cell depletion representative flow plots and absolute counts of NK1.1+ cells (per 10^6^ cells) are shown for the spleen and peripheral blood at day 8 (D). NK depletion at 3 dpi following 1e5 PFU MHV A59 infection using the timeline depicted in Figure 6A (E). Following 5e2 PFU MHV A59 infection (i.p), spleens were harvested and tittered at the indicated days post-infection with NK depletion (α-NK1.1) or non-depleting isotype control (α-Iso), with timeline depicted in Figure 6A (F). Liver histopathology scores of individual parameters (inflammation, necrosis and edema/fibrin) are shown (G). PEP-WT mice were used to verify NK cell depletion. The gating strategy to identify splenic and peripheral blood NK cells was: Lymphocytes> Single cell x2> autofluorescent-> Live> CD45+> CD3e- CD19-> NK1.1+. Representative flow plots are shown from CD3e- CD19. Each dot represents an individual mouse. p-values from Welch’s unpaired t test are shown for NK cell counts, *p<0.05, **p<0.01. Viral titers are pooled data from 2 independent experiments at each time point; comparisons are shown from multiple unpaired T-Tests, α-Iso PEP-WT vs α-NK1.1 PEP-WT , α-Iso PEP-R619W vs α-NK1.1 PEP-R619W, *p<0.01. The dashed line indicates the limit of detection (L.O.D.). For histopathological analysis, each dot represents an individual mouse, with p-values from One-way ANOVA with Tukey’s Multiple Comparisons shown, *p<0.05, **p<0.01.

**Table S1: Flow Cytometry Antibodies**

| **Antigen** | **Fluorophore** | **Dilution (1:x)** | **Company** | **Clone** | **CAT / REF** | **LOT** |
| --- | --- | --- | --- | --- | --- | --- |
| Ghost Viability | Violet510 | 1000 | TONBO Biosciences | N/A | 13-0870-T100 | D0870120221133 |
| CD45 | Spark+ BUV395 | 200 | BioLegend | 30-F11 | 103192 | B428503 |
| CD45 | BUV395 | 200 | BD Biosciences | 30-F11 | 564279 | 3205129 |
| CD3e | PECy5 | 250 | BioLegend | 145-2C11 | 100309 | B359890 |
| CD19 | BV711 | 400 | BioLegend | 6D5 | 115555 | B287454 |
| CD19 | APC | 250 | BioLegend | 6D5 | 115512 | B371440 |
| CD19 | AF700 | 250 | BioLegend | 6D5 | 115512 | B371440 |
| Ly6C | BV785 | 400 | BioLegend | HK1.4 | 128041 | B386418 |
| Ly6G | PE610 | 200 | Invitrogen | 1AB-Ly6g | 61-9668-80 | 2445867 |
| Ly6G | BV570 | 200 | BioLegend | 1AB | 127629 | B418454 |
| CD11c | PECy5.5 | 200 | Invitrogen | N418 | 35-0114-82 | 2286398 |
| CD11c | APC-Cy7 | 200 | BioLegend | N418 | 117324 | B391502 |
| CD8a | R710 | 200 | TONBO Biosciences | 53-6.7 | 80-0081-U100 | C0081010722803 |
| CD8a | AF700 | 200 | BioLegend | 53-6.7 | 100730 | B401541 |
| PDCA-1 (CD317) | Pacific Blue | 200 | BioLegend | 927 | 127108 | B302820 |
| PDCA-1 (CD317) | BV421 | 200 | BioLegend | 927 | 127023 | B365891 |
| PDCA-1 (CD317) | BV650 | 200 | BioLegend | 927 | 127019 | B365769 |
| CD80 | BV421 | 200 | BioLegend | 16-10A1 | 104725 | B366741 |
| CD86 | APC/Cy7 | 200 | BioLegend | GL-1 | 105029 | B349514 |
| CD86 | BV605 | 200 | BioLegend | GL-1 | 105037 | B373866 |
| F4/80 | Pacific Orange | 100 | Invitrogen | BM8 | MF48030 | 2209625 |
| F4/80 | PE | 200 | BioLegend | BM8 | 123109 | B223149 |
| F4/80 | FITC | 200 | BioLegend | BM8 | 123108 | B407719 |
| F4/80 | APC/Fire 810 | 200 | BioLegend | BM8 | 123166 | B431884 |
| Tim4 | PE | 150 | BioLegend | RMT4-54 | 130006 | B428998 |
| CD11b | PerCPCy5.5 | 200 | TONBO Biosciences | M1/70 | 65-0112-U100 | C0112121919653 |
| CD11b | PerCPCy5.5 | 200 | BioLegend | M1/70 | 101228 | B423267 |
| CD206 | PEcy7 | 200 | BioLegend | C068C2 | 141719 | B239603 |
| CD206 | APC | 150 | Invitrogen | MR6F3 | 17-2061-80 | 1956852 |
| MHC-II | FITC | 150 | BioLegend | M5/114.15.2 | 107606 | B398140 |
| NK1.1 | FITC | 200 | BioLegend | PK136 | 108705 | B376169 |
| NK1.1 | BV711 | 200 | BioLegend | PK136 | 108745 | B423146 |
| CD49b | FITC | 200 | eBioscience | DX5 | 11-5971-82 | E023186 |
| CD49b | PE/Dazzle 594 | 200 | BioLegend | DX5 | 108924 | B396304 |
| CD49b | PE | 200 | BioLegend | DX5 | 108908 | B391901 |
| CD49a | APC/Fire 750 | 200 | BioLegend | HM𝛼1 | 142610 | B420608 |
| CD107a | PEcy7 | 200 | BioLegend | 1D4B | 121619 | B348898 |
| CD107a | BV650 | 200 | BioLegend | 1D4B | 121645 | B424309 |
| CD27 | BV605 | 200 | BioLegend | LG.3A10 | 124249 | B408891 |
| CD27 | AF700 | 200 | BioLegend | LG.3A10 | 124240 | B329641 |
| IFNg | BV650 | 100 | BioLegend | XMG1.2 | 505832 | B361629 |
| IFNg | PE | 100 | BioLegend | XMG1.2 | 505807 | B366833 |
| Perforin | Pacific Blue | 100 | BioLegend | S160009A | 154311 | B356368 |
| Granzyme B | PE | 100 | Invitrogen | GB11 | 12-8899-41 | 2682463 |
| Granzyme B | FITC | 100 | BioLegend | GB11 | 515403 | B397296 |
| TNFa | BV605 | 100 | BioLegend | MP6-XT22 | 506329 | B422381 |
| TNFa | PE-Cy7 | 100 | BioLegend | MP6-XT22 | 506324 | B424713 |
| T-bet | BV605 | 100 | BioLegend | 4B10 | 644817 | B387805 |

List of antibodies used in various combinations throughout all experiments. Antigen, fluorophore, dilution (1:x), company, clone, catalog/reference number, and lot number listed.
